# Chromosome-level *Thlaspi arvense* genome provides new tools for translational research and for a newly domesticated cash cover crop of the cooler climates

**DOI:** 10.1101/2021.07.30.454478

**Authors:** Adam Nunn, Isaac Rodríguez-Arévalo, Zenith Tandukar, Katherine Frels, Adrián Contreras-Garrido, Pablo Carbonell-Bejerano, Panpan Zhang, Daniela Ramos-Cruz, Katharina Jandrasits, Christa Lanz, Anthony Brusa, Marie Mirouze, Kevin Dorn, Brice Jarvis, John Sedbrook, Donald L. Wyse, Christian Otto, David Langenberger, Peter F. Stadler, Detlef Weigel, M. David Marks, James A. Anderson, Claude Becker, Ratan Chopra

## Abstract

*Thlaspi arvense* (field pennycress) is being domesticated as a winter annual oilseed crop capable of improving ecosystems and intensifying agricultural productivity without increasing land use. It is a selfing diploid with a short life cycle and is amenable to genetic manipulations, making it an accessible field-based model species for genetics and epigenetics. The availability of a high quality reference genome is vital for understanding pennycress physiology and for clarifying its evolutionary history within the Brassicaceae. Here, we present a chromosome-level genome assembly of var. MN106-Ref with improved gene annotation, and use it to investigate gene structure differences between two accessions (MN108 and Spring32-10) that are highly amenable to genetic transformation. We describe small RNAs, pseudogenes, and transposable elements, and highlight tissue specific expression and methylation patterns. Resequencing of forty wild accessions provides insights into genome-wide genetic variation as well as QTL regions for flowering time and a seedling color phenotype. Altogether, these data will serve as a tool for pennycress improvement in general and for translational research across the Brassicaceae.

## Introduction

Native to Eurasia, field pennycress (*Thlaspi arvense* L.) is a member of the Brassicaceae family and is closely related to the oilseed crop species rapeseed (*Brassica rapa* and *Brassica napus* L.), camelina (*Camelina sativa* L.), as well as the wild plant *Arabidopsis thaliana* (Warwick et al., 2002; Beilstein et al., 2010). It is an emerging oil feedstock species with the potential to improve sustainability of cold climate cropping systems through use as a cash cover crop (Sedbrook et al., 2014; Chopra et al., 2018; Boateng et al., 2010). Pennycress is extremely winter hardy (Warwick et al., 2002) and can be planted in traditional fallow periods following summer annuals such as wheat, maize or soybean (Cubins et al., 2019; Johnson et al., 2015; Ott et al., 2019; Phippen and Phippen, 2012). By providing a protective living cover from the harvest of the previous summer annual crop through early spring, pennycress prevents soil erosion and nutrient loss, which in turn protects surface and below-ground water sources, suppresses early-season weed growth, and provides a food source for pollinators (Johnson et al., 2015; Weyers et al., 2021, 2019; Eberle et al., 2015). The short life cycle allows for harvest in May or June in temperate regions, with reported seed yields ranging from 750 to 2400 kg ha^-1^ (Cubins et al., 2019; Moore et al., 2020). Following harvest, an additional crop of summer annuals can be grown in a double crop system that provides increased total seed yields and beneficial ecosystem services (Phippen and Phippen, 2012; Johnson et al., 2015; Thomas et al., 2017). The pennycress seed contains an average of 30- 35% oil, and the fatty acid profile is conducive to producing biofuels (Fan et al., 2013; Moser et al., 2009; Moser, 2012). Seed oil also has the potential to be converted into an edible oil and protein source (Claver et al., 2017; Chopra et al., 2020b; McGinn et al., 2019).

*T. arvense* is a homozygous diploid species (2*n* = 2*x* = 14) (Mulligan, 1957) and is predominantly self-pollinating (Mulligan and Kevan, 1973), suggesting that breeding efforts could proceed with relative ease and speed. It is amenable to genetic transformation via the floral dip method (McGinn et al., 2019), and its diploid nature with many one-to-one gene correspondences with *A. thaliana* (Chopra et al., 2018) could provide an avenue for gene discovery followed by field-based phenotypic validation. Indeed, several agronomic and biochemical traits have already been identified in pennycress using this translational approach, including traits crucial for *de novo* domestication of *T. arvense* such as transparent testa phenotypes (Chopra et al., 2018), early flowering (Chopra et al., 2020b), reduced shatter (Chopra et al., 2020b), and seed oil composition traits (McGinn et al., 2019; Jarvis et al., 2021; Esfahanian et al., 2021; Chopra et al., 2020b). Field pennycress could thus serve as a *de novo* domesticated oilseed crop for the cooler climates of the world and at the same time as a new dicotyledonous model for functional genetics studies. Its amenability for translational research constitutes a clear advantage vis-a-vis *A. thaliana*. However, to establish *T. arvense* as a genetic model and a crop, it is important to develop genomic resources that will help explore the spectrum of genetic diversity, the extent and patterns of gene expression, genetic structure, and untapped genetic potential for crop improvement.

Here, we describe a set of new resources developed for research and breeding communities, including a high quality, chromosome-level genome assembly of *T. arvense* var. MN106-Ref, representing ∼97.5% of the estimated genome size of 539 Mbp. We provide robust annotations of both protein-coding and non-coding genes, including putative transfer RNA (tRNA), ribosomal RNA (rRNA) and small nucleolar RNA (snoRNA) predictions, alongside small RNA producing loci, transposable element (TE) families, and predicted pseudogenes. From transcriptome data based on a panel of eleven different tissues and life stages, we built a gene expression atlas. In combination with whole-genome DNA methylation profiles of both roots and shoots, this provides a basis for exploring gene regulatory and/or epigenetic mechanisms within pennycress. A comprehensive analysis of forty re-sequenced pennycress accessions highlights the nucleotide diversity in these collections, alongside gene variants and population structure. Finally, by means of genome-wide association studies (GWAS) and bulked-segregant analysis (BSA), we identified quantitative trait loci (QTL) associated with flowering time and seedling color, exemplifying the usefulness of this resource. The genome and resequencing information presented in this study will increase the value of pennycress as a model and as tool for translational research, and accelerate pennycress breeding through the discovery of genes affecting important agronomic traits.

## Results

### An improved reference genome sequence

The genome of *T. arvense* var. MN106-Ref was assembled *de novo* from 476X (256 Gb) depth PacBio Sequel II continuous long read (CLR) reads (38 kb N50). The initial assembly attempts exceeded the genome size by ∼53% with respect to the range of 459-540 Mbp total size estimated from flow cytometry and k-mer analysis **(Suppl. Table 1)**. Reducing the duplicated fraction, polishing, and scaffolding/re-scaffolding using several approaches resulted in a final assembly of ∼526 Mbp, corresponding to ∼97.5% of the upper limit of the flow cytometry-based estimate and representing an improvement of ∼20% relative to the original assembly size. Scaffolding/re-scaffolding of the genome assembly was achieved using Bionano optical, Hi-C contact, genetic linkage and comparative synteny maps. The final genome contains 964 scaffolds, with ∼83.6% of the total estimated size represented by seven large scaffolds, in agreement with the haploid chromosome number, demonstrating a vast improvement in overall contiguity and bringing the assembly to chromosome-level. The coding space is 98.7% complete on the basis of conserved core eukaryotic single-copy genes (BUSCO), with 92.1% being single-copy and 6.6% duplicated. Full descriptive statistics in comparison to T_arvense_v1 are given in **Table 1**.

**Table 1.** Full descriptive statistics comparing the previously published T_arvense_v1 assembly to the present version T_arvense_v2.

| Assembly category | T_arvense_v1 | T_arvense_v2 |
| --- | --- | --- |
| # contigs | 44,109 | 4,714 |
| Largest contig | - | 41.6 Mbp |
| contig N50 | 0.02 Mbp | 13.3 Mbp |
| # scaffolds | 6,768 | 964 |
| # scaffolds ($\geq 50,000$ bp) | 1,807 | 607 |
| Largest scaffold | 2.4 Mbp | 70.0 Mbp |
| Total length | 343 Mbp | 526 Mbp |
| Total length ($\geq 50,000$ bp) | 276 Mbp | 514 Mbp |
| GC (%) | 37.99 | 38.39 |
| N50 | 0.14 Mbp | 64.9 Mbp |
| NG50 | 0.05 Mbp | 64.9 Mbp |
| N75 | 0.06 Mbp | 61.0 Mbp |
| NG75 | - | 55.2 Mbp |
| L50 | 561 | 4 |
| LG50 | 1,678 | 4 |
| L75 | 1,469 | 6 |
| LG75 | - | 7 |
| # N's per 100 Kbp | 5,165.00 | 0.51 |

The seven largest scaffolds are all characterised by high gene density towards both telomeres and a high density of repeats and TEs in the pericentromeric and centromeric regions **(Figure 1, Suppl. Figure 1)**. Whilst the protein-coding gene fraction of the genome is similar in size to other closely-related Brassicaceae (Wang et al., 2011), the large repetitive fraction suggests an increased genome size driven by TE expansion (Beric et al., 2021). In addition, the spatial distribution of sRNA loci followed the gene density but was concentrated predominantly at the boundary between genes and TEs.

**Figure 1.**
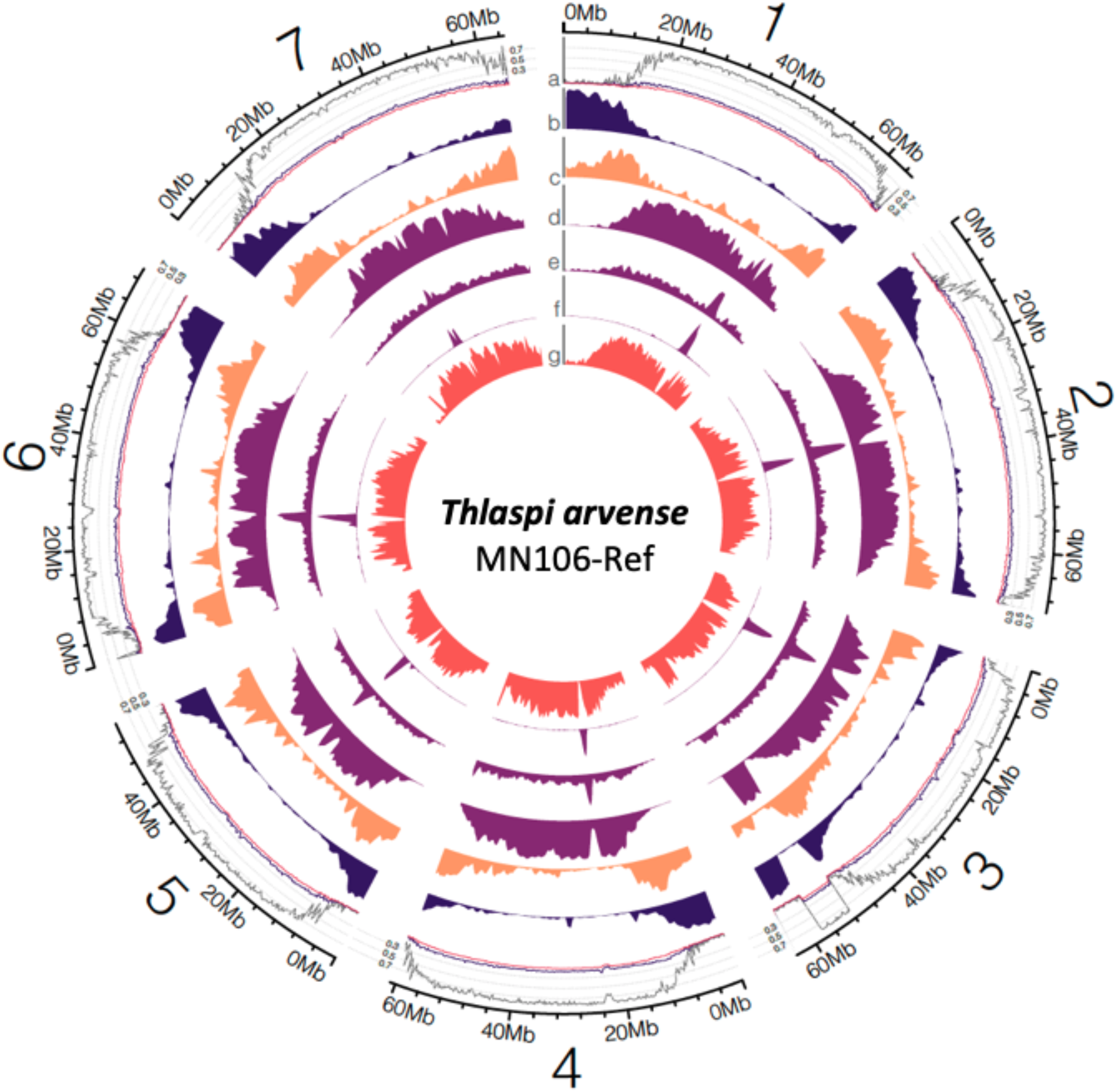
Overview of the seven largest scaffolds representing chromosomes in *T. arvense* var. MN106-Ref. The tracks denote **a)** DNA methylation level in shoot tissue (CG: grey; CHG: black; CHH: pink; 200 Kbp window size), and density distributions (1 Mbp window size) of **b)** protein-coding loci, **c)** sRNA loci, **d)** Gypsy retrotransposons, **e)** Copia retrotransposons, **f)** LTR retrotransposons, and **g)** pseudogenes.

In addition to the duplicate-containing contigs, alignments of the raw CLR reads to the new genome revealed the presence of what appeared to be a small number of collapsed repeats in scaffolds 1, 3, 5, and 7, which were typically larger than 25 Kbp and indicative of misassembly in these loci **(Suppl. Figure 2)**. Further investigation revealed an overlap with tandem repeat clusters of 18S and 28S rRNA annotations at those loci on scaffolds 3 and 5, and a large supersatellite of 5S rRNA on scaffold 1. In addition, there were corresponding genes associated with organellar DNA at those loci on scaffolds 3 and 7, indicating either erroneous incorporation of plastome sequence during assembly or genuine nuclear integrations of plastid DNA (NUPTs) (Michalovova et al., 2013).

Exploiting information from the genome of *Eutrema salsugineum* (Yang et al., 2013), a closely related species (Franzke et al., 2011) with a much smaller genome (241 Mbp) but the same karyotype (n=7), aided during re-scaffolding **(see methods; Suppl. Figure 3)** and confirmed synteny of the seven largest scaffolds in the two species **(Suppl. Figure 4)**. There is an almost 1:1 relationship between large sections of the two genomes, with the exception of some regions on scaffolds 2, 3, 6 and 7. This could be due to the low gene density observed in the *T. arvense* genome towards the center of the chromosome and/or the high presence of dispersed repeats in those regions. Chromosome evolution in the Brassicaceae has been studied through chromosome painting techniques, and 24 chromosome blocks (A-X) have been defined from an ancestral karyotype of n=8 (Schranz et al., 2006; Murat et al., 2015). We identified the 24 blocks in *T*. *arvense* based on gene homology and synteny between *T*. *arvense* and *A. thaliana* **(Figure 2)**. While in general the distribution of the chromosomal blocks resembles that in the *Eutrema* lineage (*E. salsugineum* and *S. parvula*), some blocks are rearranged in a small section at the end of the scaffold representing chromosome 1 and at the beginning of chromosome 6. The first case involves the transposition of a small part of block C in between A and B, while chromosome 6 has a possible inversion between the blocks O and W when compared to *E. salsugineum* and *S. parvula*. The synteny analysis also revealed intra-chromosomal rearrangements, but no obvious inter-chromosomal rearrangements.

**Figure 2.**
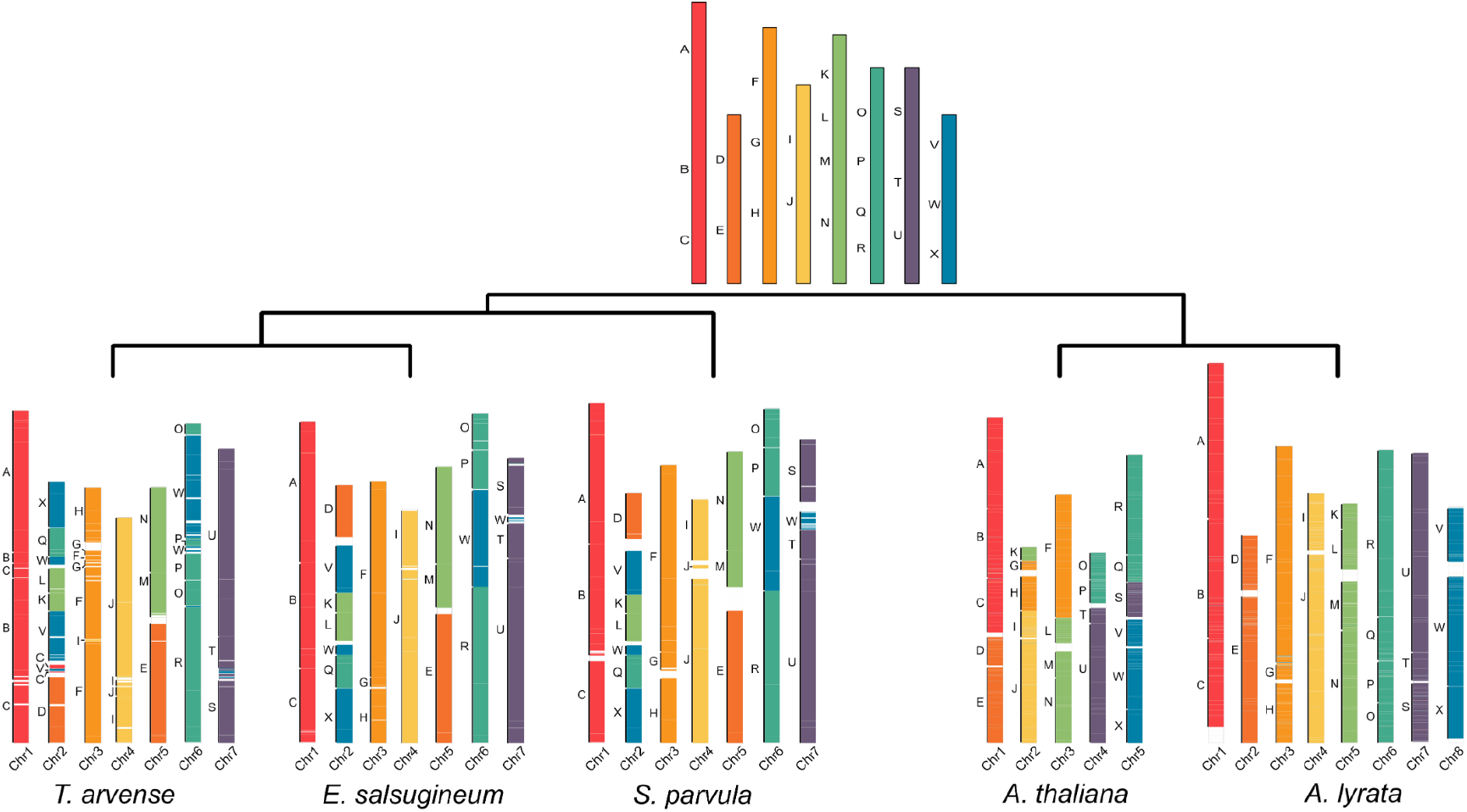
Distribution of ancestral genomic blocks (top panel) along the seven largest scaffolds of *T. arvense* MN106-Ref (T_arvense_v2), and a comparison of these genomic blocks with *Eutrema salsugineum*, *Thellungiella parvula, Arabidopsis thaliana* and *Arabidopsis lyrata*.

### Genome annotation

#### Transcriptome assembly

We sequenced total cDNA with strand-specific RNA-seq from eleven tissues, including rosette leaves, cauline leaves, inflorescences, open flowers, young green siliques, old green siliques, green seeds, mature seeds, seed pods, roots of 1-week-old seedlings, and shoots of 1-week-old seedlings **(Suppl. Table 2)**. Reads from each tissue sample were aligned to the genome with unique mapping rates between 76 and 91%, with the exception of old green silique (19%), green seed (59%), and mature seed (12%). The majority of unmapped reads in each case were due to insufficient high-quality read lengths. We constructed independent tissue-specific transcriptome assemblies and combined them into a multi-sample *de novo* assembly, yielding 30,650 consensus transcripts. These were further refined by prioritising isoforms supported by Iso-seq data, resulting in 22,124 high-quality consensus transcripts to inform gene models.

#### Protein-coding genes

In addition to the expression data, gene models were informed by protein homology using a combined database of Viridiplantae from UniProtKB/Swiss-Prot (Boutet et al., 2007) and selected Brassicaceae from RefSeq (Pruitt et al., 2012). Following initial training and annotation by *ab initio* gene predictors, protein-coding loci were further annotated with InterPro to provide PFAM domains, which were combined with a BLAST search to the UniProtKB/Swiss-Prot Viridiplantae database to infer gene ontology (GO) terms. In accordance with MAKER-P recommendations (Campbell et al., 2014), the final set of 27,128 protein-coding loci were obtained by filtering out those with an annotation edit distance (AED) score of 1 unless they also contained a PFAM domain. Approximately 95% of loci had an AED score < 0.5 **(Suppl. Figure 5)**, demonstrating a high level of support with the available evidence, and 21,171 (∼78%) were annotated with a PFAM domain. Analysis of gene orthologs and paralogs among related Brassicaceae confirmed the close relationship with *E. salsugineum*, with the protein-coding fraction occupying a genome space comparable to related species **(Figure 3a)**. A total of 4,433 gene duplication events were recorded with OrthoFinder, comparable to *E. salsugineum* (5,108), but fewer than in *B. rapa* (11,513), for example.

**Figure 3.**
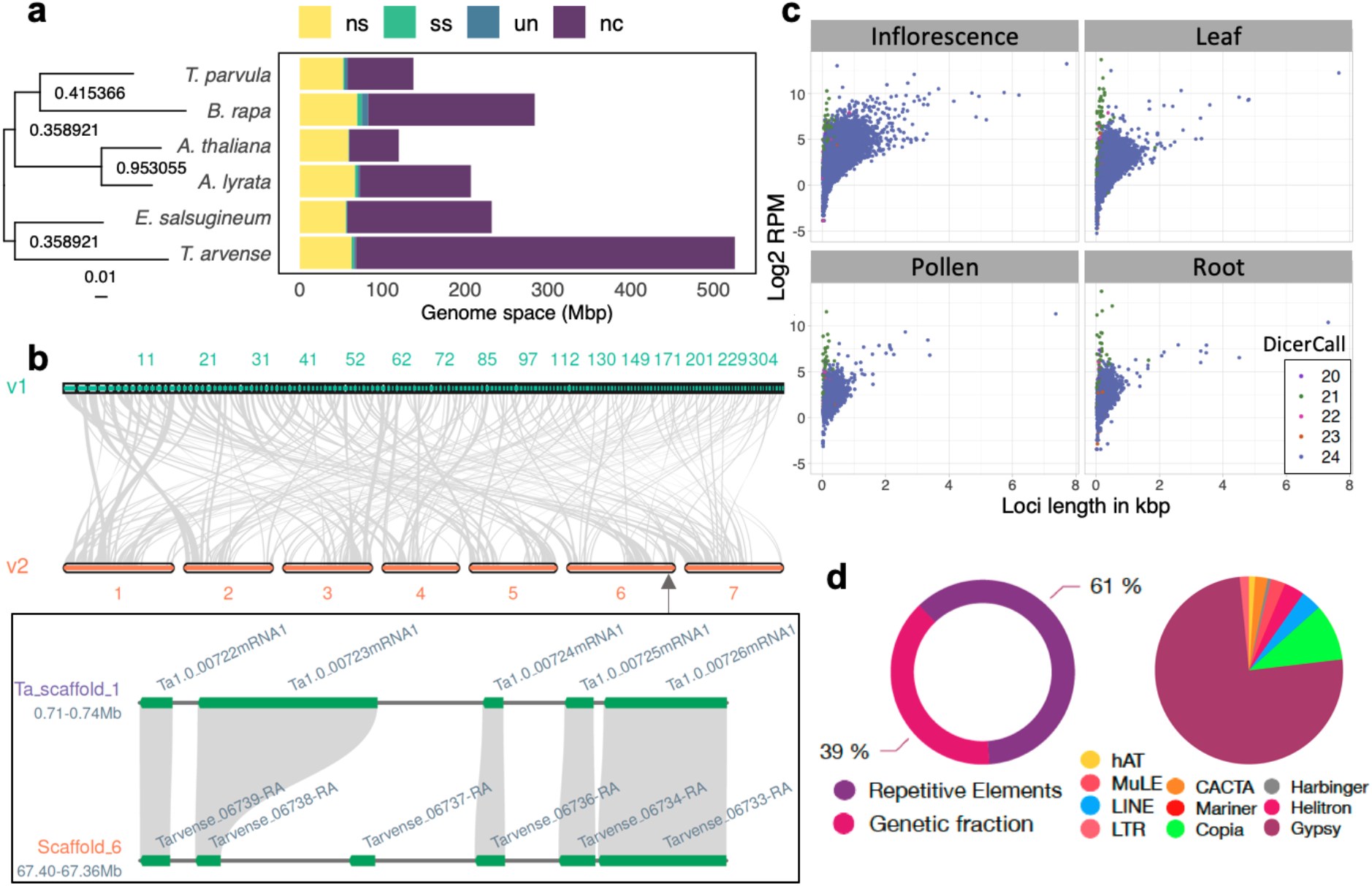
Feature annotations within *T. arvense* var MN106-Ref. **a)** Dendrogram and horizontal stacked bar chart comparing the genetic fraction in pennycress with other *Brassicaceae* sp. (ns = non-specific orthologs, ss = species-specific orthologs, un = unclassified genes, nc = non-coding/intergenic fraction). **b)** Comparison of gene macrosynteny between v1 and v2 of the genome, and a microsynteny example of genes *MYB29* and *MYB76*, which are resolved in the v2 annotation. **c)** Small RNA biogenesis loci length and expression values in each of four tissues. **d)** Overall repetitive content in the genome as discovered by RepeatMasker2, and relative abundance of TEs within the fraction of repetitive elements.

The full descriptive statistics are given in **Table 2**, in comparison to the original T_arvense_v1 annotation (Dorn et al., 2015) lifted over to the new genome with Liftoff v1.5.2 (Shumate and Salzberg, 2020), where applicable. Gene feature distributions are comparable between T_arvense_v1 and the present assembly of MN106-Ref (hereafter referred to as T_arvense_v2) (**Suppl. Figure 6**). Unique genes that were successfully lifted over from the previous version were included as a separate fraction in the final annotation (source: T_arvense_v1), resulting in 32,010 annotated genes in total. Up to ∼95.2% completeness can be obtained by combining the full set of both the current and previous annotations according to a BUSCO evaluation of 2121 conserved, single-copy orthologs. The improved contiguity of the genome space allowed for the resolution of genes such as the tandem duplicated *MYB29* and *MYB76*, which were concatenated in the previous version **(Figure 3b)**.

**Table 2.** Summary of feature annotations in comparison to the original version T_arvense_v1.

| Type | T_arvense_v1 | T_arvense_v2 | diff. |
| --- | --- | --- | --- |
| **(A) Protein-coding genes** |  |  |  |
| Total number of loci | 27,390 | 27,128 | -262 |
| Total number of unique loci | 4,780 | 5,034 | +254 |
| Total number of transcript isoforms | – | 30,650 | +30,650 |
| Number of matching loci with changes in CDS | – | – | +14,102 |
| Number of matching loci with changes in UTR(s) | – | – | +22,559 |
| Loci containing one or more PFAM domain | – | 21,171 | +21,171 |
| Loci annotated with one or more GO term | – | 13,074 | +13,074 |
| **(B) Non-coding genes** |  |  |  |
| tRNA | – | 1,148 | +1,148 |
| rRNA clusters (<25 Kbp) | – | 63 | +63 |
| snoRNA | – | 243 | +243 |
| Small interfering RNA (siRNA) | – | 19,373 | +19,373 |
| Micro RNA (miRNA) | – | 72 | +72 |
| **(C) Other gene types** |  |  |  |
| Pseudogenes (set II Ψs) | – | 44,490 | +44,490 |
| Transposable element genes | – | 423,251 | +423,251 |

#### Non-coding loci

In addition to the protein-coding gene annotations, we annotated non-coding RNA (ncRNA) genes, pseudogenes, and TEs. Descriptive annotation statistics have been summarised in **Table 2**. While many of these annotation features in *T. arvense* were similar to those found in other plant species, we observed several unique patterns, which we will describe in detail below. ncRNA annotations were inferred from sequence motifs (tRNA, rRNA, snoRNA) or from sequencing data (siRNA, miRNA). We predicted clusters of both 5S rRNA and tandem repeat units of 18S and 28S rRNA with RNAmmer (Lagesen et al., 2007), often in relative proximity to loci identified with Tandem Repeats Finder v4.09.1 (Benson, 1999) and putatively associated with centromeric repeat motifs (not shown). Of the largest seven scaffolds, only scaffolds 4 and 7 carried no such annotations. Notably, several large clusters of 5S rRNA genes were interspersed throughout the pericentromeric region of scaffold 1, whereas the remaining four scaffolds contained 18S and 28S rRNA gene annotations. Finally, we identified 243 homologs from 114 snoRNA families.

#### sRNA annotation

We identified 19,386 siRNA loci. More than 98% of these loci corresponded to heterochromatic 23-24 nt siRNA loci, with only 196 producing 20-22 nt siRNAs. The sRNA loci were expressed unevenly across tissues, as inferred from prediction with data from different tissues. Only 2,938 loci were shared across all four tissues studied (rosette leaves, roots, inflorescences, and pollen). Inflorescences were the major contributor with 6,728 private loci. Despite these differences between tissues, we observed similar overall patterns in terms of locus length, expression **(Figure 3c)**, and complexity **(Suppl. Figure 7)**.

Altogether, sRNA loci accounted for ∼8 Mbp or ∼1.5% of the assembled genome. Of the seven largest scaffolds, where the majority of genes are located, the total coverage of siRNA loci ranged between 1.5 - 2% and the loci appeared to be preferentially concentrated at the boundary between TEs and the protein-coding gene fraction of the genome. To further explore this, we partitioned the seven largest scaffolds into gene-enriched and gene-depleted regions, based on a median of 14 genes per Mbp and a mean of 54.2 genes per Mbp. We defined gene-enriched loci as those above and gene-depleted loci as those below the mean. At the chromosomal level, sRNA loci correlated with gene-enriched regions and were scarce in regions with high TE content. This trend is in contrast to that observed in *A. thaliana* (Hardcastle et al., 2018) but resembles what has been observed, for example, in maize (He et al., 2013) and tomato (Tomato Genome Consortium, 2012).

Phased secondary siRNAs (phasiRNAs) are a class of secondary sRNAs that, due to the way they are processed, produce a distinct periodical pattern of accumulation (Axtell, 2013b). In the *T. arvense* genome, we observed 139 loci with such phased patterns. In contrast to the general notion that phasiRNAs are typically 21 nt long (Lunardon et al., 2020), we found 24 nt siRNAs to be dominant in 133 of these loci.

#### MicroRNAs

MicroRNA (miRNA)-encoding genes were predicted using a combination of ShortStack and manual curation (see Methods). We identified 72 miRNA-producing loci, with 53 that were already known from other species, and 19 appeared to be species-specific. Most of the identified families were produced from only one or two loci, with miR156 and miR166 being produced by the most loci, with eight and five family members, respectively. A total of 21 out of 25 families in *T. arvense* are found in other Rosids, and three (miR161, miR157, and miR165) only in other Brassicaceae. One family, miR817, is also present in rice. There is a strong preference for 5’-U at the start of both unique and conserved miRNAs **(Suppl. Figure 8)**, in line with previous reports (Voinnet, 2009).

#### sRNA loci

When we overlaid the sRNA loci with our annotated genomic features, most sRNAs localised to the intergenic space, but a substantial fraction, especially 20-22 nt sRNAs, were produced from intronic sequences **(Suppl. Figure 9a)**. Helitrons make up only 1.5% of the genome space, yet more than 5% of sRNA biogenesis loci overlap with this type of TE. Most sRNA loci (93.0%) fell within 1.5 kbp of annotated genes or TEs **(Suppl. Figure 9b,c)**. As expected, 23-24 nt sRNAs were more frequently associated with TEs, whereas 20-22 nt sRNAs more often produced by coding genes (Axtell, 2013a).

#### Pseudogenes

In accordance with the MAKER-P protocol, pseudogenes (Ψ) were predicted in intergenic DNA with the ShiuLab pseudogene pipeline (Zou et al., 2009). A total of 44,490 set II pseudogenes were annotated, exceeding those in *A. thaliana* (∼3,700) or rice (∼7,900) by one order of magnitude. We identified 35,818 pseudogenes overlapping with TEs, and 8,672 pseudogenes that were either concentrated in intergenic space or more towards the protein-coding gene complement of the genome, and thus perhaps less likely to have arisen from retrotransposition. Approximately 59.2% of these contained neither a nonsense nor a frameshift mutation, indicating either (i) that the regulatory sequences of the pseudogenes were silenced first, (ii) a pseudo-exon which may be linked to another non-functional exon, or (iii) a possible undiscovered gene.

#### Transposable elements

In total, we identified 423,251 TEs belonging to 10 superfamilies and covering ∼61% of the genome **(Figure 3d)**. Retrotransposons (75% of all TEs are Gypsy elements; 10% Copia; 4% LINE) by far outnumbered DNA transposons (3% Helitrons; 1% hAT; 2% CACTA; 1% Pif-Harbinger; 2% MuLE). A detailed breakdown of repeats can be found in **Suppl. Table 3**. As the most abundant retrotransposon superfamily, Gypsy elements accounted for 46% of the total genome space, which is consistent with a high abundance observed in the pericentromeric heterochromatin of *E. salsugineum*, where centromere expansion is thought to have been caused by Gypsy proliferation (Zhang et al., 2020). In addition, we identified 359 protein-coding genes located fully within TE-bodies that could represent Pack-TYPE elements and contribute to gene shuffling (Catoni et al., 2018). Among these elements 153 were intersecting with mutator-like elements suggesting they correspond to Pack-MULE loci. TEs were located primarily in low gene density regions, while the fraction of TE-contained genes were randomly distributed.

### Expression atlas

With cDNA sequences from 11 different tissues or developmental stages, we could annotate tissue-specific expression patterns. The complete expression atlas is provided in **Suppl. File 1**. We evaluated the relative extent of tissue-specific gene expression using the Tau (τ) algorithm (Yanai et al., 2005), from the normalised trimmed mean of M-value (TMM) counts in all tissues (Robinson and Oshlack, 2010). To preclude potential biases caused by substantial differences in library size, we excluded low-coverage samples from mature seeds and old green siliques. In total, 4,045 genes had high or even complete tissue specificity (τ = 0.8 - 1.0), while 5,938 genes had intermediate (0.2 - 0.8) and 6,107 no or low specificity (0 - 0.2); the remaining genes were ignored due to missing data. The relative breakdown of each specificity fraction by tissue type is shown in **Figure 4a**, with “roots”, “green seeds” and “inflorescences” representing the tissues with the greatest proportion of high or complete specificity genes. The relative log2(TMM) expression values of the top 30 most highly expressed genes in each tissue, given a high or complete specificity score, are plotted in **Figure 4b** with respect to the overall mean expression per gene across all included tissues. These include, for example, genes with homology to *EXTENSIN 2* (*EXT2; A. thaliana*) in “roots”, *CRUCIFERIN* (*BnC1; B. napus*) in “green seeds”, and *PECTINESTERASE INHIBITOR 1* (*PMEI1; A. thaliana*) in “inflorescences” and “open flowers” **(Suppl. File 2)**.

**Figure 4.**
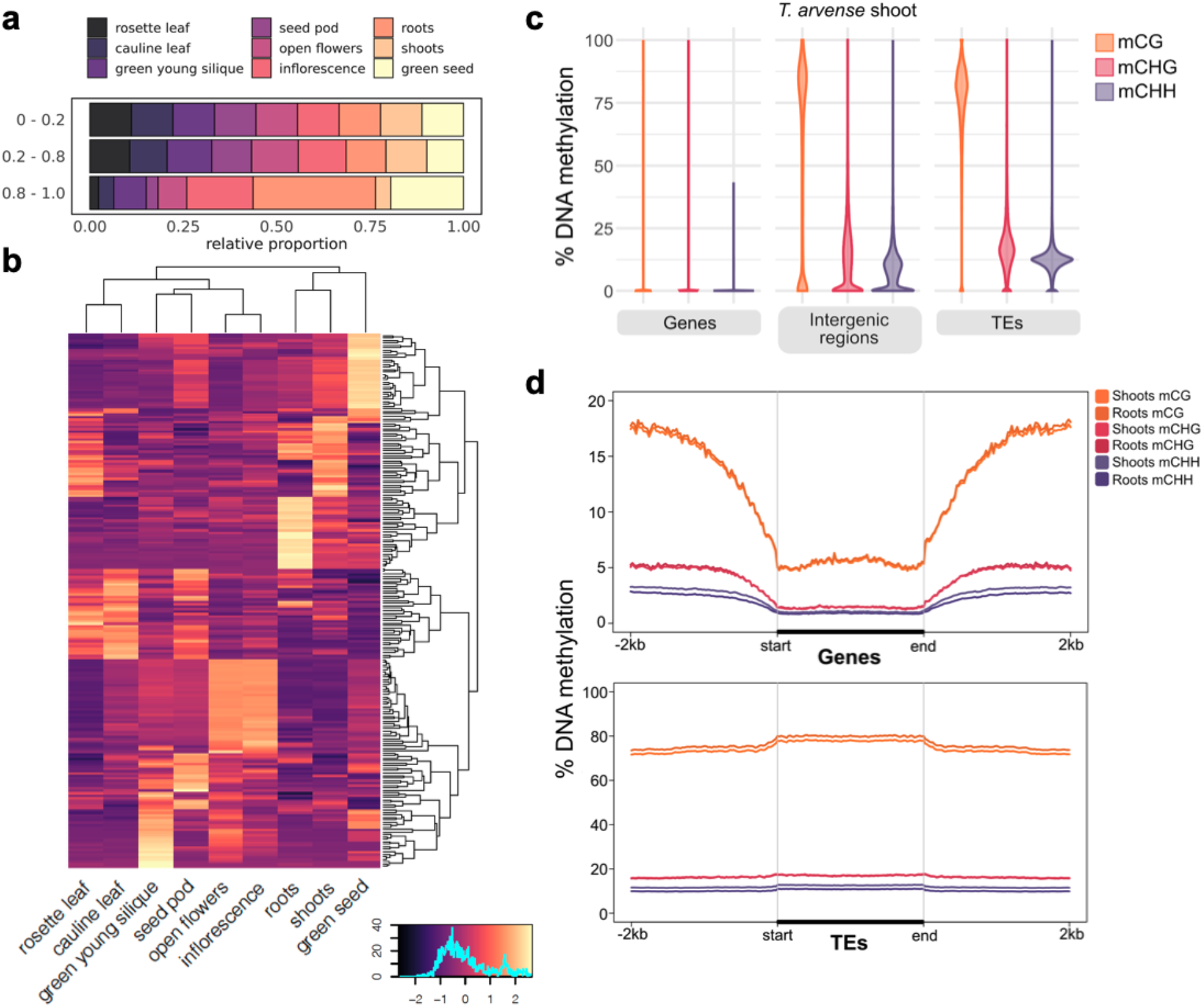
Regulatory dynamics in pennycress. **a)** Relative fraction of genes in each tissue for low (0 - 0.2), intermediate (0.2 - 0.8) and high/absolute specificity (0.8 - 1.0) subsets. **b)** Log2(TMM) expression values of the top 30 most highly expressed genes in each tissue, relative to the mean across all tissues, from the subset of genes with a high/absolute tau specificity score. **c)** Distribution of average DNA methylation for different genomic features, by cytosine sequence context. **d)** DNA methylation along genes (top) and TEs (bottom), including a 2 kb flanking sequence upstream and downstream. DNA methylation was averaged in non-overlapping 25 bp windows.

### DNA methylation

Cytosine methylation (also commonly referred to as DNA methylation) is a prevalent epigenetic mark in plant genomes and is often associated with heterochromatin and transcriptional inactivation of TEs and promoters, but also with higher and more stable expression when present in gene bodies (Zhang et al., 2018). In plants, DNA methylation occurs in three cytosine contexts, CG, CHG, and CHH (where H is any base but G), with the combined presence of CG, CHG and CHH methylation usually indicative of heterochromatin formation and TE silencing, while gene body methylation consists only of CG methylation (Bewick and Schmitz, 2017). In light of the high TE density in *T. arvense*, we analysed genome-wide DNA methylation by whole-genome bisulfite sequencing (WGBS) in shoots and roots of 2-week-old seedlings. Genome wide, 70% of cytosines were methylated in the CG context, 47% in the CHG context, and 33% in the CHH context. In line with findings in other Brassicaceae, methylation at CG sites was consistently higher than at CHG and CHH **(Figure 1a; Suppl. Figure 10)**. When we compared the WGBS data against the genome annotation, high levels of DNA methylation (mostly ^m^CG) co-localized with regions of dispersed repeats and TEs in the center of the chromosomes. Conversely, methylation was depleted in gene-rich regions **(Figure 1a,b)**. In line with this, DNA methylation was consistently high along TEs, particularly in the CG context **(Figure 4c)**. In contrast to *E. salsugineum* (Bewick et al., 2016; Niederhuth et al., 2016), DNA methylation dropped only slightly in regions flanking TEs, which might be related to the overall dense TE content in *T. arvense*.

In contrast to TE and promoter methylation, gene body methylation (gbM) is generally associated with medium-to-high gene expression levels (Zhang et al., 2006; Zilberman et al., 2006). gbM occurs in ∼30% of protein-coding genes in *A. thaliana*, with DNA methylation increasing towards the 3’-end of the gene (Zhang et al., 2006). The *T. arvense* relative *E. salsugineum* lacks gbM (Bewick et al., 2016; Niederhuth et al., 2016). gbM was also largely absent in *T. arvense* (**Figure 4d**), suggesting that gbM was lost at the base of this clade.

### Applications towards crop improvement

#### Genetic variation in a pennycress collection

Knowledge of genetic diversity within wild populations is an essential process for improvement and domestication of new crop species. We analysed a geographically broad sample of forty accessions **(Suppl. Figure 11)** using whole-genome resequencing to characterize population structure and variation in germplasm available for breeding. We identified a total of 13,224,528 variants with QD value of ≥2,000. Of these, 12,277,823 (92.8%) were SNPs, 426,115 (3.2%) were insertions, and 520,590 (3.9%) were deletions relative to the reference genome. Across all variants, 661,156 (2.9%) were in exons, with 340,132 synonymous, 314,075 nonsynonymous, and 6,949 nonsense changes. STRUCTURE analysis of both indel and SNP datasets resulted in optimal models of k=3 populations **(Suppl. Figure 12)**. Both data sets assigned the three lines of Armenian descent, which were highly distinct and had the largest genetic distance to the other accessions, to a single discrete population with limited to no gene flow to the other populations. These results are consistent with previous reports in pennycress (Frels et al., 2019) and were further supported by whole-genome dendrograms **(Figure 5a)**. We also calculated linkage disequilibrium (LD) among 2,213,286 genome-wide markers and chromosome-specific markers using TASSEL v5.2.72 (Bradbury et al., 2007) with a sliding window of 40 markers. The r-squared values were plotted against the physical distance with a LOESS curve fitted to the data to show LD decay **(Suppl. Figure 13)**. Genome-wide, LD decayed to an r-squared value (*r^2^*) of 0.2 over 6 kbp (Hill and Weir, 1988), which is comparable to LD decay reported in related Brassica species at *r^2^* = 0.3, including *B*. *rapa* (2.1 Kbp) (Wu et al., 2019)and *B*. *napus* (12.1Kbp) (Lu et al., 2019).

**Figure 5.**
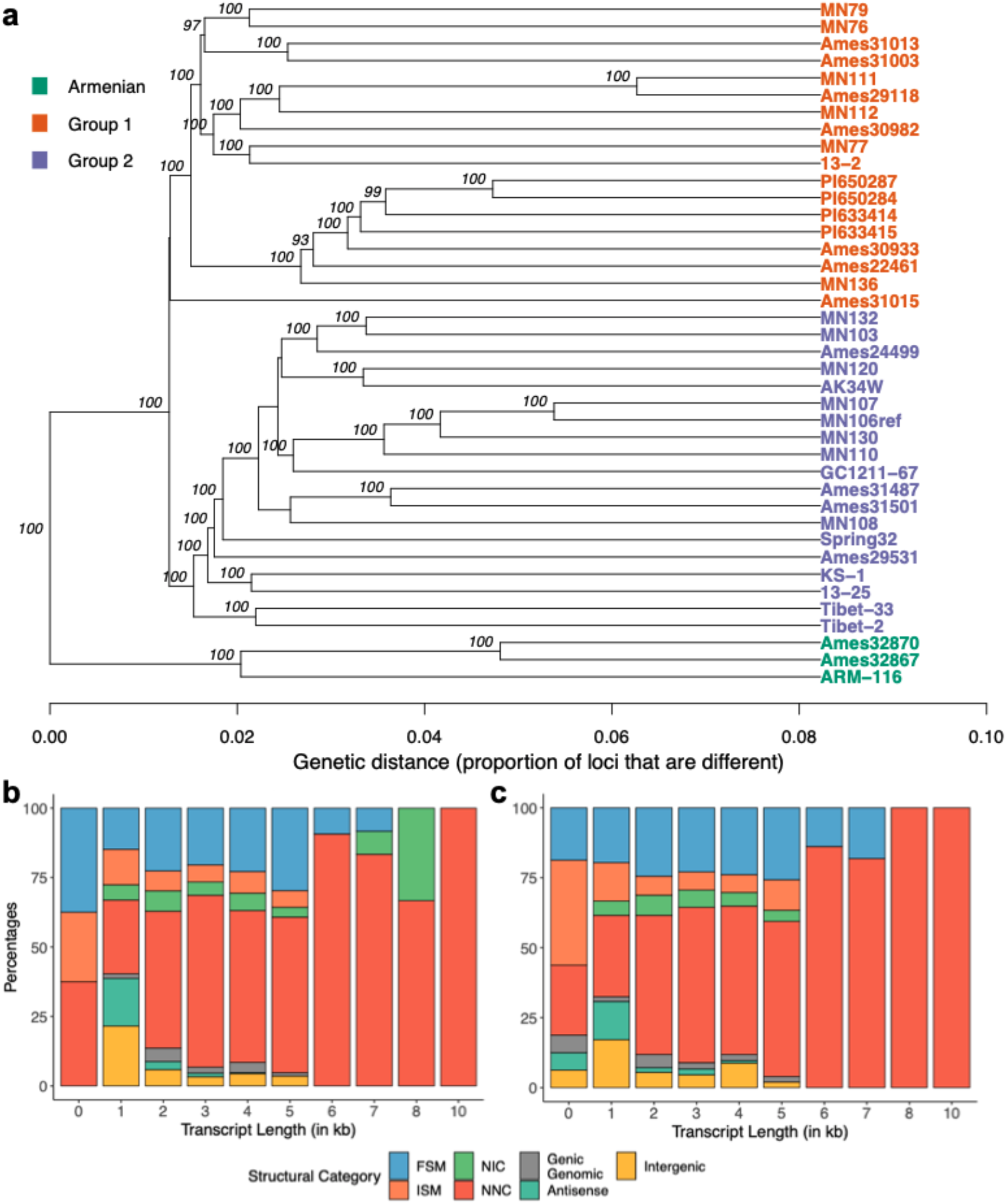
a) Dendrogram representing the forty wild accessions in our study showing three distinct sub- populations, inferred from STRUCTURE analysis (Suppl. Figure 11). **b,c)** Variation of transcript isoforms for MN108 **(b)**, and Spring32-10 **(c)** accessions based on SQANTI3 analysis.

#### Gene structure variation in pennycress accessions

The natural variation present in germplasm is an important source of alleles to facilitate breeding efforts and presents an opportunity to understand the evolution of gene families and adaptation within a species. To understand these in a more targeted approach, we sequenced on the PacBio Sequel platform the transcriptomes of two accessions, MN108 and Spring32-10, that are amenable to transformation and gene editing (McGinn et al., 2019), using RNA from leaves, roots, seeds, flowers and siliques. We constructed *de novo* reference transcriptomes using the Iso-seq3 pipeline, resulting in 25,296 and 26,571 accession-specific isoforms for MN108 and Spring32-10, respectively. These transcriptomes were then polished using the raw reads and processed through the SQANTI3 pipeline (Tardaguila et al., 2018) to characterize the genes and isoforms identified in each of the accessions. We identified 212/220 unique genes and 3,780/3,857 unique isoforms for MN108/Spring32-10 respectively compared to the new reference. Transcripts mapping to the known reference denoted by “Full Splice Match” (FSM) and “Incomplete Splice Match” (ISM) accounted for 28.7% and 30.6% of al transcript models in MN108 and Spring32-10, respectively **(Figure 5b,c)**. Transcripts of the antisense, intergenic, and genic intron categories collectively accounted for a total of 12.0% (MN108) and 11.2% (Spring32-10). About ∼15% of all identified transcripts were novel isoforms when compared to the reference transcriptome for T_arvense_v2.

#### Genome-wide association analysis for flowering time in pennycress collections

For pennycress to fit between the rotations of traditional summer crops, it needs to mature in the spring without interfering with the planting of the following summer crop. Hence, it becomes important to understand the genetic control of flowering time and to identify natural alleles responsible for this trait in field pennycress. We performed a genome-wide association (GWA) analysis using whole genome resequencing data filtered at a minor allele frequency of 5%, leaving 534,758 SNPs, and a Bonferroni corrected genome-wide significance threshold. The analysis detected two significant SNPs, one each on scaffold 3 and 6, associated with flowering time in climate controlled conditions **(Figure 6a)**. The SNP marker with the highest significant association was located ∼1.3Mbp from the *FLC* locus on scaffold 6 **(Figure 6a)**, a well-known flowering regulator that confers a winter annual habit in Brassicaceae and other species (Tadege et al., 2001; Schiessl et al., 2019).

**Figure 6.**
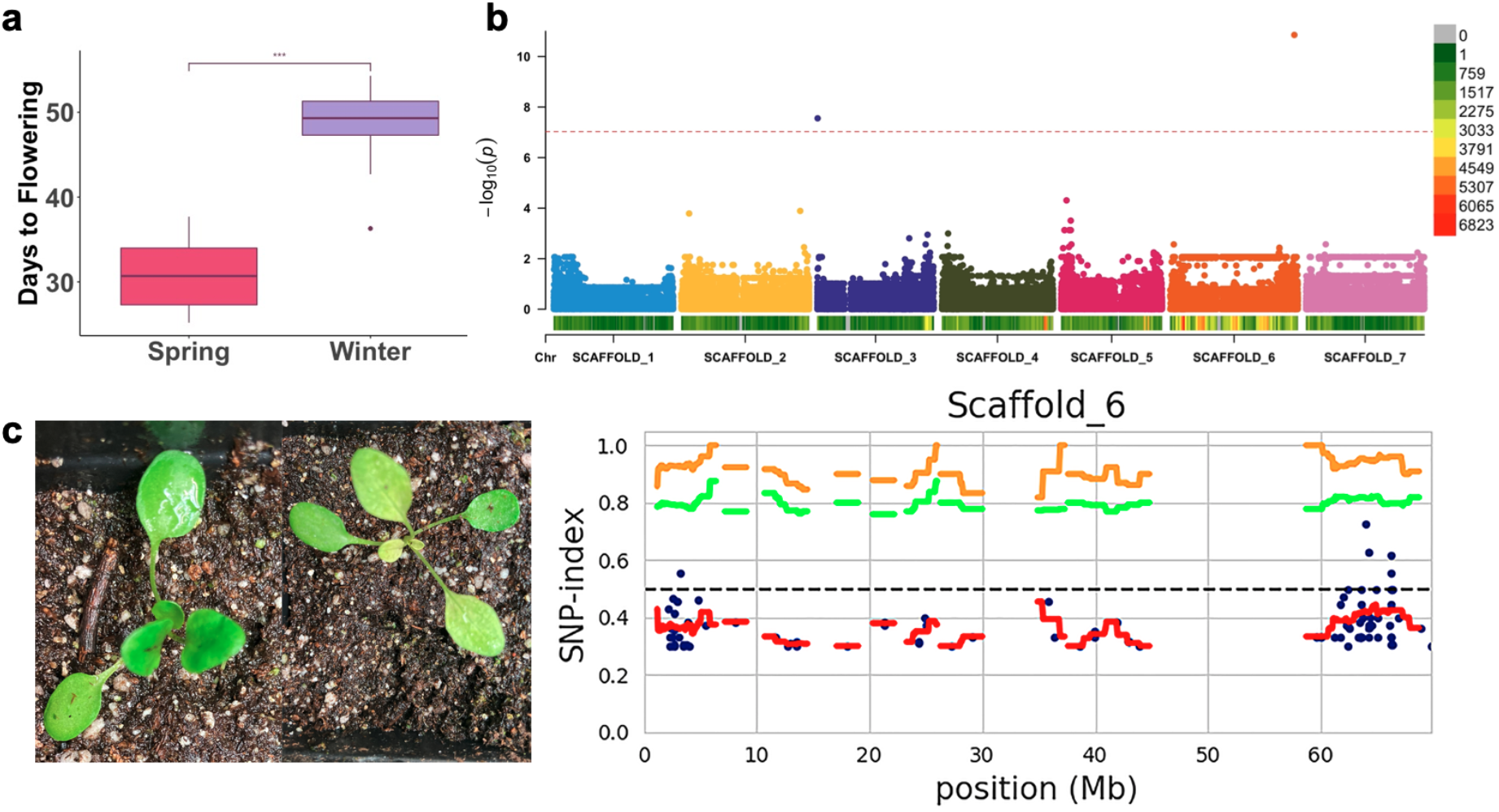
Mapping studies were performed using the improved pennycress genome assembly. **a)** Days to flowering in forty resequenced accessions representing a collection from various regions of the world (Suppl. Figure 11). **b)** Manhattan plot highlighting QTLs associated with this trait. **c)** A pale phenotype segregating in an improved pennycress line (*fae1-1*/*rod1-1*) was analyzed with a modified bulked segregant analysis and the QTL region associated with this phenotype was mapped using the MutMap approach.

#### Mapping a pale seedling phenotype

From a segregating population with a high oleic pennycress (*fae-1/rod1-1*) background (Chopra et al., 2020b), we identified pale seedling lines **(Figure 6b)**. This phenotype segregated in a Mendelian fashion. To determine the genetic control for this phenotype, we separately pooled genomic DNA from 20 wild-type and 20 pale plants. We processed sequence data obtained from each of these pools through the MutMap pipeline (Sugihara et al., 2020) and discovered a putative genomic interval (63.85 - 63.95 Mbp) on scaffold 6 linked to the pale phenotype. SnpEff (Cingolani et al., 2012) identified polymorphisms that might have deleterious effects on function of genes in this region **(Suppl. Table 4)**. The most obvious candidate is *MEX1*, encoding a maltose transporter located in the chloroplast, knockout of which causes a pale seedling phenotype in *A. thaliana* (Niittylä et al., 2004).

## Discussion

In this study, we report a high-quality reference genome assembly and annotation for *T. arvense* (var. MN106-Ref), a newly domesticated oilseed crop for the cooler climates of the world. The improved genome assembly, containing seven chromosome-level scaffolds, revealed two main features: a landscape characterised by a large repetitive fraction populated with TEs and pseudogenic loci in pericentromeric regions, and a gene complement similar in size to other Brassicaceae and densely concentrated towards the telomeres **(Figure 1)**. Previous annotations were enriched with additional gene models for protein-coding loci, and now include non-coding genes for tRNAs, rRNAs, snoRNAs, siRNAs and miRNAs, alongside predicted pseudogenes and TEs **(Table 2)**. These newly improved assembly features will allow for efficient combining of traits and help accelerate future breeding as it would provide knowledge about the gene localization and the linkage of genes of interest. For example, the improved genome assembly has revealed that multiple domestication syndrome genes (*ALKENYL HYDROXALKYL PRODUCING 2 - like, Transparent Testa 8, EARLY FLOWERING 6*) **(Suppl. Figure 1)** are located on a single chromosome.

Improved genomic resources can facilitate general understanding of plant biology and evolutionary biology while aiding plant breeding and crop improvement (Scheben et al., 2016). For example, pennycress and *Arabidopsis* share many key features that made *Arabidopsis* the most widely studied model plant system (Meinke et al., 1998). The use of *Arabidopsis* for translational research and for identifying potential gene targets in *T. arvense* is possible and has been extensively validated (Chopra et al., 2018, 2020b; McGinn et al., 2019; Jarvis et al., 2021; Chopra et al., 2020a). Previous studies have suggested that over a thousand unique genes in *T. arvense* are represented by multiple genes in *Arabidopsis* and vice versa. Our comparative genomics by way of synteny with *E. salsugineum* (Yang et al., 2013) revealed a high level of agreement, particularly between the protein-coding fraction of the genome, represented as conserved blocks in the largest seven scaffolds relative to the ancestral karyotype in Brassicaceae (Murat et al., 2015) **(Figure 2)**. The detailed description of gene synteny between *T. arvense* and other Brassicaceae provides insights into the evolutionary relevance of *T. arvense* within lineage II of Brassicaceae. In addition, the difference in genome size between *T*. *arvense* and other species, despite the reduced level of gene duplication and the 1:1 gene relationship, can be explained by the large repetitive fractions present throughout both the centromeric and pericentromeric regions. In the absence of whole genome duplication events, these repetitive fractions indicate that the increased genome size may be a consequence of active TE expansion. This is therefore suggestive of a mechanism by which deleterious retrotransposon insertions must be mitigated in *T. arvense*. This could be explained by the high proportion of *Gyspy* retrotransposons in this species, usually located in heterochromatic regions, or by integration site selection (Sultana et al., 2017), or otherwise via silencing by small RNA activity and/or DNA methylation (Bucher et al., 2012; Sigman and Slotkin, 2016). Given the relatively high error rate of PacBio CLR reads (∼10% before correction) with respect to circular consensus sequencing (CCS), the repetitive fraction would also help to explain the initial overestimation of the assembly size as a result of duplicated contigs. We also detected several loci with highly overrepresented read coverage indicative of repeat collapsing during the assembly process, often intersecting with 5S, 18S, and 28S rRNA annotations. Such regions are difficult even for current long read technologies due to the large size of the tandem repeat units.

With the availability of improved genomic resources, increasing interest has turned towards understanding tissue-specific gene regulation to reduce pleiotropic effects upon direct targeting of genes during crop improvement. In this study, we have generated a resource using mRNA-seq, sRNA-seq and WGBS to gain insights into genes and their associated regulatory landscape. These datasets help elucidate the extent of tissue specificity and provide useful information for gene modification targets. For example, fatty-acid desaturase 2 gene (FAD2; *Ta12495* - T_arvense_v1) is involved in the oil biosynthesis pathway and is expressed in many different tissues analyzed in this study **(Suppl. File 1)**. *FAD2* gene knockout should result in higher levels of oleic acid in the seed oil and provide an opportunity for pennycress oil to be used in food applications. It has been observed, however, that knockout mutants in pennycress display delayed growth and reduced seed yields in spring-types (Jarvis et al., 2021), and reduced winter survival in the winter-types (Chopra et al., 2019), as a purported consequence of its broad expression profile. Similarly, genes such as *AOP2-LIKE* (*Tarvense_05380* - T_arvense_v2) have been targeted to reduce glucosinolates in pennycress seed meal for food and animal feed applications (Chopra et al., 2020b). However, *AOP2-LIKE*, too, is expressed in many tissues during development, which might explain why knockout plants with reduced glucosinolate content are reportedly more susceptible to insect herbivores such as flea beetles feeding on rosette leaves and root tissues (Marks et al., 2021). Our tissue-specific expression data suggest that, to overcome this challenge, one could alternatively target genes such as *Glucosinolate Transporter 1* (GTR1; *Tarvense_14683*), which is expressed specifically in reproductive tissues **(Suppl. File 1).** This might achieve the desired reductions of seed glucosinolates while avoiding developmental defects. Such approaches have been effectively used in *Arabidopsis* and many *Brassica* species (Nour-Eldin et al., 2012; Andersen and Halkier, 2014).

Finally, the forty resequenced accessions described here provide a rich source of variants that reflect the genetic diversity and population structure of the species in the collection (**Figure 5a**). Further evaluations of transcriptome sequences showed ample variation in the transcripts from two separate lines - MN108 and Spring 32-10 - that are highly amenable to transformation and highlighted the potential for developing pan-genomes in the future. These genomic resources will facilitate genetic mapping studies in pennycress in both natural populations and mutant panels. We have identified genomic regions associated with variation in flowering time using GWAS methods (**Figure 6a**) and a pale leaf phenotype in pennycress seedlings using a modified BSA-Seq approach (**Figure 6b**).

Over the last few years, significant efforts have been made towards the discovery of crucial traits and translational research in pennycress, centering on MN106-Ref and the gene space information generated by Dorn et al. (2013 and 2015). In this study we continued to generate genomic tools for this accession, with improved contiguity and high quality annotations to make *T. arvense* var. MN106-Ref more accessible as a field-based model species for genetics and epigenetics studies and to provide tools for this new and extremely hardy winter annual cash cover crop. However, the assembly of additional accessions can only help to further enrich the resources available for the study of pennycress. In parallel to this study, a Chinese accession of *T. arvense* (YUN_Tarv_1.0) was assembled using Oxford Nanopore, Illumina HiSeq and Hi-C sequencing (Geng et al., 2021). This timely availability of an additional frame of reference opens the door to a pan-genomic approach in evolutionary research, and allows for the better characterisation of structural variants moving forward. Furthermore, the use of different sequencing technologies and assembly software provides an additional avenue to correct mis- assemblies and base calling errors in either case. The overall longer contigs assembled with PacBio CLR, for example, and the consideration of various genetic map data in addition to Hi-C, provides a greater resolution of scaffolds particularly throughout the centromere and pericentromeric regions (Suppl. Figure 16). The reduced error-rate of PacBio CCS (used for polishing) is also reflected in the overall k-mer content, which is measured with a two-order magnitude higher consensus quality over scaffolds representing chromosomes and ∼99% overall completeness for T_arvense_v2 (Suppl. Tables 7-9), indicative of high-quality, error-free sequences more appropriate for variant calling, for instance. Geng et al. (2021) also reported WGS analysis on forty Chinese accessions and reported an LD decay of 150 kbp at an r-squared value (*r^2^*) of 0.6, which is considerably higher than the values determined on the forty accessions in this study, as well as those reported for related *Brassica* species (Wu et al., 2019; Lu et al., 2019). We believe the combination of resources will allow us to investigate the differences that might exist between accessions originating from different geographic locations around the world, and help provide further insight into structural variations and evolutionary dynamics.

In conclusion, the T_arvense_v2 assembly offers new insights into the genome structure of this species in particular and of lineage II of Brassicacae more generally, and it provides new information and resources relevant for comparative genomic studies. The tools presented here provide a solid foundation for future studies in an alternative model species and an emerging crop.

## Methods

### Seeds for the reference genome development

Seeds from a small natural population of *T. arvense* L. were collected near Coates, MN by Dr. Wyse and the accession number MN106 was assigned to this population. We propagated a single plant for ten generations from this population and we refer to this line as MN106-Ref.

### Sample collection, library preparation, and DNA sequencing for assembly

#### PacBio CLR library

Plants were cultivated, sampled and prepared at the Max Planck Institute for Developmental Biology (Tübingen, Germany). Plant seeds were stratified in the dark at 4°C for 4-6 d prior to planting on soil. Samples were collected from young rosette leaves of *T. arvense* var. MN106-Ref seedlings, cultivated for two weeks under growth chamber conditions of 16-23°C, 65% relative humidity, and a light-dark photoperiod of 16 h:8 h under 110-140 μmolm^-2^s^-1^ light. High molecular weight (HMW) DNA was obtained following nuclei isolation and DNA extraction with the Circulomics Nanobind Plant Nuclei Big DNA kit according to the protocol described in Workman et al. (2018) (Workman et al., 2019). A total of 11 extractions from 1.5-2 g frozen leaves each were processed in that way, yielding a pooled sample with a total of 12 μg of DNA by Qubit estimation, and high DNA purity with a mean absorbance ratio of 1.81 at 260/280 nm absorbance and 2.00 at 260/230 nm absorbance, as measured by Nanodrop spectrophotometer. HMW DNA was sheared by one pass through a 26G needle using a 1 mL syringe, resulting in an 85 kb peak size sample as estimated by Femto Pulse Analyzer. A large insert gDNA library for PacBio Sequel II CLR sequencing was prepared using the SMRTbell® Express Template Preparation Kit 2.0. The library was size-selected for >30 kb using BluePippin with a 0.75% agarose cassette (Sage Science) and loaded into one Sequel II SMRT cell at a 32 pM concentration. This yielded a genome-wide sequencing depth of approximately 476X over ∼6.9 million polymerase reads with a subread N50 of ∼38 kbp.

#### PacBio CCS library

MN106-Ref plants were grown in growth chambers at the University of Minnesota. Individual plants were grown to form large rosettes for isolating DNA. Approximately 25 g of tissue was harvested and submitted to Intact Genomics (Saint Louis, MO, USA) for High Molecular Weight DNA extraction. This yielded a pooled sample with a total of 269 ng of DNA by Qubit estimation, and high DNA purity with a mean absorbance ratio of 1.87 at 260/280 nm absorbance and 2.37 at 260/230 nm absorbance, as measured by Nanodrop spectrophotometer. To further clean up the high molecular weight DNA, we used Salt:Chloroform Wash protocol recommended by PacBio (Shared-Protocol-Guidelines-for-Using-a-Salt-Chloroform-Wash-to-Clean-Up-gDNA.pdf). This yielded a total of 12.1 ng/ul of high quality DNA for library preparation. A large insert gDNA library was prepared and 15 kb High Pass Size Selection on Pippin HT was performed at the University of Minnesota Genomics Center (Minneapolis, MN, USA). These libraries were sequenced on 4 SMRT cells using PacBio Sequel II.

#### Bionano library

High-molecular-weight DNA was isolated from young leaves and nicking endonuclease - BspQI was chosen to label high-quality HMW DNA molecules. The nicked DNA molecules were then stained as previously described (Lam et al., 2012). The stained and labelled DNA samples were loaded onto the NanoChannel array (Bionano Genomics, San Diego, CA, USA) and automatically imaged by the Irys system (Bionano Genomics, San Diego, CA, USA).

#### Hi-C library

The MN106-Ref plant tissue used for PacBio CCS was submitted to Phase Genomics (San Diego, CA, USA). The Hi-C library was prepared following the proximo Hi-C plant protocol (Phase Genomics, San Diego, CA, USA) and the libraries were sequenced to 116X depth on an Illumina platform with the paired-end mode and read length of 150 bp.

#### Illumina PCR-free library

Libraries for PCR-free short read sequencing were prepared from MN106-Ref genomic DNA using the TruSeq DNA PCR-Free Low Throughput Library Prep Kit (Illumina, San Diego, CA, USA) in combination with TruSeq DNA Single Indexes Set A (Illumina, San Diego, CA, USA) according to the manufacturer’s protocol. We prepared two libraries, with average insert sizes of 350 bp and 550 bp, respectively. Samples were sequenced to 125X depth (∼66 Gb) on an Illumina HiSeq2500 instrument with 125 bp paired-end reads.

#### Genome assembly and construction of chromosome-level scaffolds

The initial assembly was performed using Canu v1.9 (Koren et al., 2017) with default options, aside from cluster runtime configuration and the settings `corOutCoverage=50`, `minReadLength=5000`, `minOverlapLength=4000`, `correctedErrorRate=0.04` and `genomeSize=539m`, which were selected based on the characteristics of the library. Canu performs consensus-based read correction and trimming, resulting in a curated set of reads which were taken forward for assembly (**Suppl. Figure 14**).

The resulting assembly overestimated the genome size by approximately 53%, which we surmised was likely due to uncorrected sequencing errors in the remaining fraction of reads which Canu was able to assemble into independent, duplicated contigs. Analysis of single-copy orthologs from the *Eudicotyledons odb10* database with BUSCO v3.0.2 (Simão et al., 2015) revealed a high completeness of 98.4% and a duplication level of 23.6% (**Suppl. Table 5**). Subsequent alignment of the reads to the assembly using minimap2 v2.17 (Li, 2018) and purge_dups v1.0.1 (Guan et al., 2020) presented bimodal peaks in the read depth distribution, indicative of a large duplicated fraction within the assembly (**Suppl. Figure 15**). As efforts to collapse this duplicated fraction using assembly parameters were unsuccessful, and purge_dups is intended to correct duplication arising from heterozygosity (which does not apply in *T. arvense*), the fraction was reduced by manual curation instead. Contigs starting from the left-hand side of the read depth distribution were consecutively removed until reaching an approximation of the estimated genome size, with any contigs containing non-duplicated predicted BUSCO genes kept preferentially in favour of discarding the next contig with lower read depth in the series.

The deduplicated assembly from Canu was polished with the PacBio Sequel II HiFi CCS reads using two iterations of RACON v1.4.3 (Vaser et al., 2017), prior to repeat re-assembly. Bionano maps were used to build *de novo* scaffolds using the polished assembly; hybrid scaffolds were generated using the *de novo* Bionano maps and the assembly (https://bionanogenomics.com/support-page/data-analysis-documentation/). To further resolve repetitive regions and improve assembly contiguity, the bionano-scaffolded assembly was integrated into the HERA pipeline (Du and Liang, 2019). The Hi-C data was aligned with bwa mem v0.7.17 (Li and Durbin, 2009), PCR duplicates marked with picard tools v1.83 (http://broadinstitute.github.io/picard) and the quality assessed with the `hic_qc.py` tool of Phase Genomics (https://github.com/phasegenomics/hic_qc). The assembly was then scaffolded with the Hi-C alignments using SALSA v2.2 (Ghurye et al., 2017) and subsequently polished with the PCR-free Illumina data using two iterations of PILON v1.23 (Walker et al., 2014). The final assembly was the result of a meta-assembly with quickmerge v0.3 (Chakraborty et al., 2016), which combined the current assembly with an earlier draft version assembled using Canu 1.8 (Koren et al., 2017) directly from the PacBio CCS reads and polished only with the Illumina PCR-free short-reads, following an almost identical workflow, in order to help address the possibility of misassembly arising from technical sources and improve overall contiguity. This resulting assembly was evaluated with BUSCO (Simão et al., 2015) and QUAST v5.0.2 (Gurevich et al., 2013).

#### Genome size estimation using flow cytometry and k-mer based approach

The nuclei of field pennycress line MN106-Ref, *Arabidopsis thaliana*, Maize, and Tomato were stained, with propidium iodide and fluorescent signals were captured using a flow cytometer. DNA content for all four species that corresponded to G0/1 nuclei are listed in **Suppl. Table 1**. The genome size of Arabidopsis is 135 Mb and therefore, the genome size of pennycress was calculated to be 501± 33 Mb. Using the Illumina HiSeq2500 platform, we obtained ∼100x PCR-free reads, which were used for subsequent K-mer analysis using Jellyfish (Marçais and Kingsford, 2011). The 101-mer frequency distribution curve exhibited a peak at 22-kmer and analysis showed that the total number of K-mers was 11,403,836,319. Using the formula of genome size = total K-mer number/peak depth, the genome size of this sequencing sample was estimated to be 518,356,196 bp. Similarly, the single copy content of the genome was estimated to reach 79%. Using both methods of genome size estimation, we found the pennycress genome ranged from 459 to 540 Mb.

#### Development of genetic maps for re-scaffolding

To improve the contiguity and correct misassemblies, we developed two genetic linkage maps using F2 populations. The first linkage map was derived from a cross between a wild Minnesota accession “MN106-Ref’’ and a genetically distant Armenian accession “Ames32867”. The resulting F1 plants were allowed to self-fertilize, and seeds from a single plant were collected and propagated to the F2 generation. Approximately 500 mg fresh tissue was collected from 94 individuals in the F2 population. The tissue was desiccated using silica beads and pulverized using a tissue lyser. DNA was isolated with the Biosprint DNA plant kit (Qiagen, Valencia, CA, USA). The F2 population along with the two parental genotypes was genotyped with genotyping-by-sequencing at the University of Minnesota Genomics Center (Minneapolis, MN, USA). Each sample was digested with the *BtgI*_*BgLII* restriction enzyme combination, barcoded, and sequenced on the Illumina NovaSeq S1 (Single-end 101 bp) yielding 1,237,890 mean reads per sample. The raw reads were de-multiplexed based on the barcode information and aligned to the most recent iteration of the pennycress genome using bwa. Sequence aligned files were processed through samtools v1.9 (Li et al., 2009) and picard tools to sort the files and remove group identifiers. Variants were called using GATK HaplotypeCaller v3.3.0. SNPs identified among these 94 lines were used for the development of genetic maps. The second linkage map was derived from a cross between MN106-Ref and a mutant line “2019-M2-111’’. To identify the variant alleles in 2019-M2-111, we performed whole genome re-sequencing using paired-end reads on the Illumina Platform. SNPs were identified using a similar approach as described above. Sixty-seven SNP markers were designed using the bi-allelic information from resequence data. DNA was extracted from 48 samples from the mutant F2 population using the Sigma–Aldrich ready extract method, allele-specific and flanking primers synthesized from IDT (Iowa, USA) for each of the alleles were mixed (**Suppl. File 3**), and genotyping was performed using the methods described in (Chopra et al., 2020a).

A total of 35,436 SNPs were identified among the population used for the first linkage map, SNP sites were selected with no-missing data, QD > 1000 and the segregation of the markers was 1:2:1. A total of 743 high quality SNPs were retained for further analysis. A genetic map for the population was constructed using JoinMap 5 (Stam, 1993). Only biallelic SNPs were used in the analysis and genetic maps were constructed with regression mapping based on default parameters of recombination frequency of <0.4 with only the first two steps. The Kosambi mapping function was chosen for map distance estimation, and the Ripple function was deployed to confirm marker order within each of the seven linkage groups. A total of 319 markers were mapped to seven linkage groups **Suppl. File 4**. Similarly, 67 markers were genotyped on 48 individuals from the second population of linkage and 52 markers were mapped to six linkage groups **Suppl. File 5**. Both of these linkage maps were used reordering and correcting the scaffolds as described below.

#### Re-scaffolding

Initial exploration regarding gene and TE distributions and methylation patterns pointed to potential mis-assemblies in the assembled genome. Further investigation by way of synteny comparison to a closely-related species, *Eutrema salsugineum* (Yang et al., 2013), revealed that several of these likely occurred during scaffolding as orientation errors. Consequently, we manually introduced breakpoints at selected loci in the assembled genome where they were supported by at least two sources of data from synteny maps (derived from reciprocal best blast), genetic linkage maps (wild-derived and EMS mutation based) and Hi-C. These were cross-examined with minimap2 alignments of PacBio CLR reads to the genome, an overview of corresponding gene distributions produced by Liftoff v1.5.2 (Shumate and Salzberg, 2020), and the resulting synteny analysis to *E. salsugineum*. The resulting contigs were then re-scaffolded with ALLMAPS v1.1.5 (Tang et al., 2015) to produce the final assembly, integrating both the synteny map and genetic map data and manually discounting contigs that were supported only by single markers.

### Genome annotation

#### Tissue preparation for RNA sequencing

*T. arvense* var. MN106-Ref seeds were surface sterilized with chlorine gas for 1 h and stratified for 3 d at 4°C. For seedling-stage RNA extractions, seeds were plated on ½ MS medium supplement with 1% plant Agar and stratified for 3 d at 4°C. For all other tissue collections, plants were sown on soil and grown in a climate-controlled growth chamber in long-day conditions (16/8 h light/dark at 21°/16°C, light intensity 140 µE / m^2^*s, with 60% relative humidity; plants were watered twice per week. Two weeks after germination, plants growing on soil were vernalized at 4°C in the dark for 4 weeks, then moved back to the growth chamber. Samples were collected from 11 different tissues in three biological replicates (two in case of mature seeds); for each replicate, we pooled tissue from two individuals. Tissues included: one-week old shoots (from plate culture), one-week old roots (from plate culture), rosette leaves, cauline leaves, inflorescences, open flowers, young green siliques (about 0.5×0.5 cm), older green siliques (about 1×1 cm), seed pods, green seeds, and mature seeds.

#### RNA extraction and sequencing

Total mRNA was extracted using the RNeasy Plant Kit (Qiagen, Valencia, CA, USA) and treated with DNase I using the DNA-free kit DNase Treatment and Removal Reagents (Ambion by life technologies, Carlsbad, CA, USA), following the manufacturer’s protocols. cDNA libraries were constructed using the NEBNext Ultra II Directional RNA Library Prep kit (New England BioLabs, Ipswich, MA, USA inc.) for Illumina following the manufacturer’s protocol. Libraries were sequenced on a HiSeq 2500 instrument (Illumina, San Diego, CA, USA) as 125 bp paired-end reads.

#### Transcriptome assembly

Following quality control and adapter clipping with cutadapt (Martin, 2011), biological replicates for each of eleven tissue types from Illumina mRNA-seq libraries were aligned independently using STAR v2.5.3a (Dobin et al., 2013), then merged according to tissue type, prior to assembly via a reference-based approach. Each assembly was performed using Ryuto v1.3m (Gatter and Stadler, 2019), and consensus reconstruction was then performed using TACO v0.7.3 (Niknafs et al., 2017) to merge tissue-specific transcriptome assemblies. PacBio Iso-seq libraries from MN106-Ref were refined, clustered and polished following the Iso-seq3 pipeline (https://github.com/PacificBiosciences/IsoSeq), prior to alignment with STARlong and isoform collapsing using the cDNA_Cupcake (https://github.com/Magdoll/cDNA_Cupcake) suite. The Iso-seq data was later leveraged together with the Illumina mRNA-seq data to prioritise convergent isoforms using custom in-house scripting.

#### Genome annotation

The final assembly was annotated using the MAKER-P v2.31.10 (Campbell et al., 2013, 2014) pipeline on the servers provided by the EpiDiverse project, at ecSeq Bioinformatics GmbH (Leipzig, Germany). Plant proteins were obtained from the *Viridiplantae* fraction of UniProtKB/Swiss-Prot and combined with RefSeq sequences derived from selected Brassicaceae: *Arabidopsis thaliana*, *Brassica napus*, *Brassica rapa*, *Camelina sativa* and *Raphanus sativus*. TEs were obtained from RepetDB (Amselem et al., 2019) for selected plant species: *Arabidopsis lyrata*, *Arabidopsis thaliana*, *Arabis alpina*, *Brassica rapa*, *Capsella rubella*, and *Thellungiella parvula (Eutrema parvulum)*. Repeat library construction was carried out using RepeatModeler v1.0.11 (Smit and Hubley, 2008) following basic recommendations from MAKER-P (Campbell et al., 2014). Putative gene fragments were filtered out following BLAST search to the combined Swiss-Prot + RefSeq protein plant database after exclusion of hits from RepetDB. The *de novo* library was combined with a manually curated library of plant sequences derived from repbase (Bao et al., 2015). Genome masking is performed with RepeatMasker v4.0.9 (Smit, 2004) as part of the MAKER-P pipeline. Protein-coding genes, non-coding RNAs and pseudogenes were annotated with the MAKER-P pipeline following two iterative rounds under default settings, using (i) transcript isoforms from Illumina mRNA-seq and PacBio Iso-seq data, (ii) protein homology evidence from the custom Swiss-Prot + RefSeq plant protein database, and (iii) the repeat library and TE sequences for masking. The initial results were used to train gene models for *ab initio* predictors SNAP v2006-07-28 (Korf, 2004) and Augustus v3.3.3 (Stanke et al., 2006), which were fed back into the pipeline for the subsequent rounds. The final set of annotations were filtered based on Annotation Edit Distance (AED) < 1 except in cases with corresponding PFAM domains, as derived from InterProScan v5.45-80.0 (Jones et al., 2014). The tRNA annotation was performed with tRNAscan-SE v1.3.1 (Lowe and Eddy, 1997) and the rRNA annotation with RNAmmer v1.2 (Lagesen et al., 2007), respectively. The snoRNA homologs were derived using Infernal v1.1.4 (Nawrocki and Eddy, 2013) from plant snoRNA families described in (Patra Bhattacharya et al., 2016). Gene orthologs and duplication events in comparison to selected Brassicaceae (*A. lyrata*, *A. thaliana, B. rapa*, *E. parvulum*, and *E. salsugineum*) were evaluated with OrthoFinder v2.5.2.

#### Transposable element annotation

Two *de novo* annotation tools, EDTA v1.7.0 (Ou et al., 2019) and RepeatModeler v2.0 (Flynn et al., 2020), were used to annotate TEs independently. For EDTA the following parameters were used in addition to defaults: `--species others`, `--step all`, `--sensitive 1`, `--anno 1`, and `--evaluate 1`. For RepeatModeler2 the additional parameters were `-engine ncbi` and `-LTRStruct`. The outputs of both tools were evaluated by manual curation. First, we used tblastn to align each TE consensus with the transposase database obtained from repbase, and the retrotransposon domains (GAG, Pol, Env, etc.) were viewed one by one with dotter (Sonnhammer and Durbin, 1995). Sequences with multiple paralogs were mapped back to the genome and manually extended to determine the full length boundary of each TE. A total of 107 full length, representative *Copia* and *Gypsy* families were successfully evaluated. The TE consensus from RepeatModeler2 was selected as the most accurate model based on full length paralogs. RepeatMasker was then used to construct the GFF3- like file from the FASTA file from RepeatModeler2, with the optional settings: `-e ncbi -q - no_is -norna -nolow -div 40 -cutoff 225`. The perl script `rmOutToGFF3.pl` was used to generate the final GFF3 file.

#### sRNA plant material

Seeds were sterilized by overnight incubation at −80°C, followed by 4 hours of bleach treatment at room temperature (seeds in open 2 mL tube in a desiccator containing a beaker with 40 mL chlorine-based bleach (<5%) and 1 mL HCl (32%)). For rosette, inflorescence and pollen, seeds were stratified in the dark at 4°C for six days prior to planting on soil, then cultivated under growth chamber conditions of 16-23°C, 65% relative humidity, and a light-dark photoperiod of 16h:8h under 110-140 μmol m^-2^ s^-1^ light. Rosette leaves were harvested after two weeks of growth. For inflorescence and pollen, six week old plants were vernalized for four weeks at 4°C in a light-dark photoperiod of 12h:12h under 110-140 μmol m^-2^ s^-1^ light. Two weeks after bolting, inflorescence and pollen was collected. Pollen grains were collected by vortexing open flowers in 18% sucrose for 5 min followed by centrifugation at 3,000g for 3 min in a swinging bucket rotor. For root samples, seeds were stratified for six days at 4°C in the dark on ½ MS media. Plants were grown in 3-4 mL ½ MS medium plates in long-day (16 hours) at 16°C. Root samples were collected 12-14 days after stratification.

#### sRNA extraction and library preparation

Total RNA was extracted by freezing collected samples with liquid nitrogen and grinding with a mortar and pestle with Trizol reagent (Life Technologies, Carlsbad, CA, USA). Then, total RNA (1 μg) was treated with DNase I (Thermo Fisher Scientific, Waltham, MA, USA) and used for library preparation. Small RNA libraries were prepared as indicated by the TruSeq small RNA library prep kit (Illumina, San Diego, CA, USA), using 1 μg of total RNA as input, as described by the TruSeq RNA sample prep V2 guide (Illumina, San Diego, CA, USA). Size selection was performed using the BluePippin System (SAGE Science, Massachusetts, USA). Single-end sequencing was performed on a HiSeq3000 instrument (Illumina, San Diego, CA, USA).

#### sRNA loci annotation

Raw FASTQ files were processed to remove the 3′-adapter and quality controlled with trim_galore v0.6.6 (https://www.bioinformatics.babraham.ac.uk/projects/trim_galore/) using `trim_galore –q 30 --small_rna`. Read quality was checked with FastQC v0.11.9 (https://www.bioinformatics.babraham.ac.uk/projects/fastqc/). The reference annotation of sRNA loci was created following the steps indicated by (Lunardon et al., 2020). In short, each library was aligned to the reference genome independently using ShortStack v3.8.5 (Axtell, 2013b), with default parameters, to identify clusters of sRNAs *de novo* with a minimum expression threshold of 2 reads per million (RPM). sRNA clusters from all libraries of the same tissue were intersected using BEDTools v2.26.0 `multiIntersectBed` (Quinlan and Hall, 2010) with default parameters, and only those loci present in at least three libraries were retained. For each tissue, sRNA clusters 25 nt apart were padded together with the `bedtools merge -d` option. sRNA loci whose expression was <0.5 RPM in all libraries of each tissue were also removed. Finally, sRNA loci for all different tissues were merged in a single file retaining tissue of origin information with `bedtools merge -o distinct` options. miRNAs predicted by the ShortStack tool were manually curated (**Suppl. Inf. 1**) following the criteria of (Axtell, 2013b): Maximum hairpin length of 300 nt; ≥ 75% of reads mapping to the hairpin must belong to the miRNA/miRNA* duplex; For the miRNA/miRNA* duplex: No internal loops allowed, two-nucleotide 3′ overhangs, maximum five mismatched bases, only three of which are nucleotides in asymmetric bulges; Mature miRNA sequence should be between 20 nt and 24 nt.

### Expression atlas

Gene expression was measured from the same tissue-specific STAR alignments taken prior to merging biological replicates for transcript assembly, excluding coverage outliers “mature seed” and “green old silique”. A total of 27 samples from 9 tissues were therefore considered for gene expression analysis. Raw counts were generated using subread featureCounts v2.0.1 (Liao et al., 2014) and subsequently normalised using the trimmed mean of M-values (TMM) (Robinson and Oshlack, 2010) derived from edgeR v3.34 (Robinson et al., 2010). Averaged expression counts by group were taken for tissue specificity evaluation using the Tau (τ) algorithm (Yanai et al., 2005), as implemented in the R package tispec v0.99.0 (https://rdrr.io/github/roonysgalbi/tispec/), which provides a measure of τ in the range of 0 - 1 where 0 is non/low specificity and 1 indicates high/absolute specificity.

### DNA methylation

We extracted genomic DNA from roots and shoots of 2-week-old seedlings grown on ½ MS medium with 0.8% Agar and 0.1% DMSO. Seedlings were grown vertically in 16h/8h light/dark; at the time of sampling, roots were separated from shoot tissue with a razor blade and the plant tissue was flash-frozen in liquid nitrogen. Genomic DNA was extracted from ground tissue using the DNeasy Plant Mini kit (QIAGEN, Hilden, Germany). Libraries for WGBS were prepared using the NEBNext UltraII DNA Library Prep kit (New England Biolabs). Adapter-ligated DNA was treated with sodium bisulfite using the EpiTect Plus Bisulfite kit (QIAGEN, Hilden, Germany) and amplified using the Kapa HiFi Uracil + ReadyMix (Roche, Basel, Switzerland) in 10 PCR cycles. WGBS libraries were sequenced on an Illumina HiSeq2500 instrument with 125 bp paired-end reads.

The WGBS libraries were processed using the nf-core/methylseq v1.5 pipeline (<u>10.5281/zenodo.2555454</u>) combining bwa-meth v0.2.2 (Pedersen et al., 2014) as an aligner and MethylDackel v0.5.0 (https://github.com/dpryan79/MethylDackel) for the methylation calling. The default parameters were used for the entire workflow with the exception of the methylation calling where the following arguments were used: `-D 1000 --maxVariantFrac 0.4 -- minOppositeDepth 5 --CHG --CHH --nOT 3,3,3,3 --nOB 3,3,3,3 -d 3`. Only cytosines with a minimum coverage of 3x were kept for the subsequent analysis. Further comparisons between the methylated cytosines and the genome annotation were performed using BEDtools v2.27.1 (Quinlan and Hall, 2010).

### Population genomics

DNA from forty pennycress accessions was extracted from approximately 500 mg of leaf tissue pooled from five plants using a plant genomic DNA kit (Epoch Life Science). DNA was then subjected to whole genome sequencing on an Illumina Novaseq sequencer (2 x 125 bp). Raw reads were then aligned to the new reference genome (T_arvense_v2) using bwa-mem (Li and Durbin, 2009). The aligned files were processed with Samtools and Picard tools and variants were called using GATK HaplotypeCaller v3.3.0 (Ren et al., 2018). Variants were annotated using SnpEff 5.0e (Cingolani et al., 2012). Datasets for both Indel and SNP panels were trimmed based on LD prior to population genomic analysis using Plink v1.9 (Purcell et al., 2007) with the parameter `--indep-pairwise 1000 5 0.5`. Population structure for both SNP and indel data were then characterized using the admixture model and independent allele frequencies in STRUCTURE v2.3.4 (Pritchard et al., 2000). Dendrograms of both SNP and Indel data were generated under the UPGMA method using the R package poppr (Kamvar et al., 2014).

#### Structural variants using Iso-seq data

Single-molecule real-time (SMRT) isoform sequencing (Iso-seq) based on PacBio (Pacific Biosciences, Menlo Park, USA) generated reads were used to investigate unambiguous full length isoforms for two pennycress wild accessions, MN108 and Spring32-10. Total RNA extraction was performed on the green seed, hypocotyl, seedling root, and flower tissues from pennycress plants grown in a climate controlled growth chamber maintained 21°C/20°C during 16h:8h day-night setting. Approximately 250 ng of total RNA was obtained and subjected to the Iso-seq Express Library Workflow (Pacific Biosciences, Menlo Park, CA, USA). cDNA is synthesized from full length mRNA with the NEBNext Single Cell/ Low Input RNA prep kit followed by PCR amplification. The amplified cDNA is converted into SMRTbell templates using the PacBio SMRTbell Express Template Prep Kit 2.0 for sequencing on the Sequel System. Sequencing was performed at the University of Minnesota Genomics Center facility (Minneapolis, MN, USA).

The polished high quality FASTA file obtained from Iso-seq3 was aligned to pennycress version 2 (T_arvense_v2) with minimap2 (Li, 2018). The resulting SAM file was sorted and collapsed using the cDNA_Cupcake package to obtain an input GFF file such that each transcript has exactly one alignment and at most one ORF prediction. `Sqanti3_qc.py`, part of the SQANTI3 package (Tardaguila et al., 2018), was deployed on the resulting GFF file along with the reference genome in the FASTA format and a GTF annotation file. This returned a reference corrected transcriptome, transcript-level and junction-level files with structural and quality descriptors, and a QC graphical report. Among the splice junction sites, SQANTI3 defines canonical junctions as AT-AC, GC-AG, GT-AG, whereas all others are classified as non-canonical splice junctions.

#### Linkage disequilibrium analysis

Linkage disequilibrium (LD) among genome-wide markers and chromosome-specific markers were calculated with TASSEL v5.2.72 (Bradbury et al., 2007) with a sliding window size of 40 markers with 88,530,900 total comparisons. The r-squared values obtained via the linkage disequilibrium function in TASSEL were plotted against the physical distance with a LOESS curve fitted to the data to show LD decay **(Suppl. Figure 13)**.

#### Genome-wide association analysis

The forty accessions were planted in a three replication, randomized complete block design in a greenhouse maintained at 21°C/20°C and 16 hour days. Ten seeds per replicate were planted in 13.3 cm^2^ pots in Sungrow propagation potting mix. Seedlings were thinned to one plant per pot after emergence. Winter annual accessions require vernalization to induce flowering, so all winter accessions were placed in a growth chamber maintained at 4 °C with 16 h light for a period of 21 days about 4 weeks after emergence. Spring annual accessions were planted approximately five weeks after winter accessions. Data for days to flowering were collected on 34 accessions that germinated as the number of days that elapsed from the date of emergence to the appearance of the first flower. The vernalization requirement for winter accessions explains the large differences in mean number of days to flowering between spring and winter accessions. Additional phenotypes and data associated with these sequenced accessions is available in **Suppl. File 6.**

The multi-locus model farmCPU (Liu et al., 2016) in the rMVP package (Yin et al., 2021) for R statistical software was used to perform a genome wide association study for flowering time among 34 wild accessions. The “farmCPU” model was preferred to other models as it improves statistical power, computationally efficiency, prevents model overfitting, and controls for false positives by incorporating population structure and kinship in a fixed and random mixed linear model. Three principal components were added as covariates in the analysis to account for population structure that exists within the germplasm. A total of 534,758 biallelic SNPs were used in the analysis with the genome-wide significance threshold determined by Bonferroni correction (ɑ/n) where ɑ is set at 0.05 and n is the total number of SNPs used.

### Bulked-Segregation sequencing and MutMap analysis

Bulked Segregant Analysis (BSA) (Michelmore et al., 1991) coupled with whole genome sequencing (BSA-Seq) was performed to locate the genomic region harboring the gene responsible for the pale mutant phenotype in pennycress **(Figure 6b)**. Two pools were created with one pool containing leaf tissue from 20 individual pale mutants and the other pool consisting of wild-type individuals that did not exhibit the pale phenotype. DNA was extracted from fresh pennycress leaves using the DNeasy Plant Mini Kit (QIAGEN, Valencia, CA, USA). Both pools were sequenced on an Illumina HiSeq 2000 instrument using 2 x 125 base paired reads at the University of Minnesota Genomics Center (Minneapolis, MN, USA). The reads were analyzed using the MutMap pipeline (Sugihara et al., 2020) and the QTL region was surveyed for the candidate genes.

### Comparison to YUN_Tarv_1.0

Synteny between T_arvense_v2 and YUN_Tarv_1.0 was assessed with minimap2 alignments and the resulting dotplot generated with the R package dotPlotly (https://github.com/tpoorten/dotPlotly). The k-mer analysis of quality and completeness was carried out for each assembly with Merqury v1.3 (Rhie et al., 2020), using both the PCR-free Illumina HiSeq reads generated in this study and those obtained from Geng et al. (2021) under the accession SRR14757813 in the NCBI Sequence Read Archive.

## Data availability

The assembly and all NGS-based raw data are deposited in the ENA Sequence Read Archive repository (www.ebi.ac.uk/ena/) under study accession number PRJEB46635. Summary of data provided by each institute and corresponding application has been described in **Suppl. Table 6.**

## Acknowledgements

We thank David Galbraith, Win Phippen, Thomas Gatter, MPI DB Genome Center, Vienna Biocenter Core Facilities (VBCF), University of Minnesota Genomics Center (UMGC), Minnesota Supercomputing Institute (MSI), and all EpiDiverse network members and beneficiaries. We acknowledge the hard work of many who contributed to this study including Brett Heim, Krishan Rai, Nicole Folstad, Matthew A. Ott, Shweta Jain, and many others.

## Author contributions

RC, AN, KF, and CB conceived the study. RC and AN led the genome assembly and evaluation, assisted by IRA, PCB. IRA performed the comparative genomics analysis of synteny during genome re-scaffolding and in the final evaluation. AN led the genome annotation and performed analysis for protein-coding genes, non-coding genes (tRNA, rRNA, snoRNA), pseudogenes. Small RNA library, annotation and analysis was performed by ACG, supervised by DW. PZ and ACG performed the transposable element annotation, supervised by DW and MM. AN performed the gene expression analysis and evaluation of tissue specificity. CB and KJ provided PCR-free libraries and RC performed k-mer analysis for genome estimation. KF and RC provided the CCS libraries and was prepared by UMGC. CB and IRA provided the DNA methylation libraries and analysis. RC, ZT, MDM, and KF developed linkage mapping populations, designed primers, performed genotyping and built genetic maps. KF, RC, and KD generated resources for HiC, Bionano and resequencing of accessions. KF and ZT phenotyped resequenced accessions. RC performed SNP analysis of resequenced datasets. ZT performed the linkage disequilibrium decay analysis. AB performed population genomics and ZT and RC performed GWAS analysis. RC and BJ prepared samples for Iso-seq libraries and RC and ZT performed gene structure variation analysis. RC and MDM performed bulk-segregant analysis. The PacBio CLR library was prepared and sequenced by PCB and AN under the guidance of CL. DR prepared and sequenced mRNA-seq libraries. The manuscript was written by RC, AN, CB, ACG, IRA, and ZT and reviewed by all authors. All authors have approved the manuscript.

## Supplementary Legends

**Supplementary Figure 1.** Karyotype plot of the seven largest scaffolds representing chromosomes in T. arvense MN106-Ref (T_arvense_v2), alongside a concatenation of all minor scaffolds. Transposable element LTR annotations (red) are highlighted as an approximate localisation of each centromere, alongside loci containing putative telomeric repeat motifs (black labels) and genes of interest (blue labels) in the *de novo* domestication of pennycress. Scaling is given in Mbp.

**Supplementary Figure 2.** Integrative Genome Viewer (IGV) snapshot of PacBio read coverage (top track) over the largest seven scaffolds of the genome, including distributions of genes (middle track) and transposable elements (bottom track). Spikes in coverage in scaffolds 1, 3, 5, and 7 are indicative of collapsed repeats, which are typically larger than the average read length.

**Supplementary Figure 3.** Sequence dot plots showing the largest seven scaffolds of the closely-related species *E. salsugineum* and their equivalent in *T. arvense* var. MN106-Ref (T_arvense_v2), comparing the difference both **a)** before and **b)** after re-scaffolding. Horizontal dashed lines in (a) denote breakpoints which were manually introduced to the genome based on evaluation of genetic maps, synteny maps and Hi-C data.

**Supplementary Figure 4.** Synteny analysis between the largest seven scaffolds of the closely-related species *E. salsugineum* and their equivalent in *T. arvense* var. MN106-Ref (T_arvense_v2). Ribbons show the syntenic relationships between the two genomes. Dark ribbons indicate syntenic blocks in inverse orientation.

**Supplementary Figure 5.** The cumulative distribution of annotation edit distance (AED) scores from the final set of protein-coding loci, denoting that ∼95% of annotated genes are supported with a score ≤ 0.5 overall.

**Supplementary Figure 6.** An overview of annotated genomic feature distributions in comparison to T_arvense_v1 for **a)** gene lengths, **b)** CDS lengths, **c)** per gene exon number, and **d)** intron lengths.

**Supplementary Figure 7.** Small RNA (sRNA) annotation in the T_arvense_v2 genome assembly. **a)** sRNA loci per tissue of origin (RPM = reads per million). **b)** sRNA complexity, measured as “number of distinct alignments / total number of alignments”. **c)** sRNA locus size distribution. **d)** Co-occurrence of sRNAs between tissues. Colored horizontal bars show the total number of loci per tissue.

**Supplementary Figure 8.** MiRNAs in the T_arvense_v2 genome assembly. **a)** Size of mature miRNAs identified in this study, split into novel and conserved miRNA species. **b-d)** Sequence conservation in miRNAs, measured in bits foreach position of the mature microRNA (Schneider and Stephens 1990). Sequences were aligned from the 5’-end. Different panels show sequence conservation of all miRNAs **(b)**, only novel miRNAs **(c)** and only conserved miRNAs **(d)**.

**Supplementary Figure 9.** sRNA types and their association with different genomic features. **a)** Occupancy of all annotated siRNAs and miRNAs in either genes, TE superfamilies, or intergenic regions. **b,c)** Association of sRNA loci with either TEs or genes within 1.5 Kb distance, for all sRNAs **(b)** and for only phased loci **(c)**.

**Supplementary Figure 10.** Methylation rate frequency distribution by sequence context in shoot and root tissues.

**Supplementary Figure 11.** Map showing original sampling sites of pennycress accessions used for resequencing analysis in this study.

**Supplementary Figure 12.** Structure plot showing inferred population membership for SNP data (top) and Indel data (bottom) at k = 3 for the re-sequenced accessions.

**Supplementary Figure 13.** Genome-wide linkage disequilibrium decay plotted against physical distance for MN106-Ref (T_arvense_v2) at an r-squared value of 0.2 and chromosome level LD decay described in the right. Linkage disequilibrium (LD) was calculated using 2,213,286 genome-wide markers with a sliding window of 40 markers.

**Supplementary Figure 14.** Read length distribution of trimmed PacBio Sequel II HiFi CLR reads taken forward for assembly with Canu v1.9.

**Supplementary Figure 15.** Distribution of PacBio Sequel II HiFi CLR read mapping depth frequency over assembled contigs, with bimodal peaks due to contig regions with lower depth than the average indicating that they are duplicated.

**Supplementary Figure 16.** Synteny between T_arvense_v2 (x-axis) and YUN_Tarv_1.0 (y-axis). Significantly more un-scaffolded contigs from YUN_Tarv_1.0 map to T_arvense_v2 than vice versa, with a total of 44 query sequences from the Chinese accession retained after post-filtering for minimum alignment length 5,000 and aggregate alignment length of 500,000. Most un-scaffolded contigs map to pericentromeric and centromeric regions, which are visible here in addition to the notable mis-scaffolding of centromeric repeats at the telomeric ends of the chromosome-representing scaffolds in YUN_Tarv_1.0.

**Supplementary Table 1.** Estimation of the genome size of *T. arvense* using flow cytometry with *Arabidopsis thaliana*, tomato (*Solanum lycopersicum*), and maize (*Zea mays*) as references.

**Supplementary Table 2.** Alignment statistics of mRNA-seq reads prior to merging by tissue type.

**Supplementary Table 3.** Detailed per-class statistics of the transposable element fraction of the *T. arvense* genome.

**Supplementary Table 4.** Description of genes identified in the QTL region (Scaffold_6: 63.85 - 63.95 Mbp) of the BSA analysis of pale seedling phenotype in pennycress.

**Supplementary Table 5.** BUSCO statistics on **a)** initial assembly, immediately after CANU, and **b)** final assembly. Both are derived from orthologs to the *Eudicotyledons odb10* database.

**Supplementary Table 6.** Summary of data provided by each institute and corresponding application.

**Supplementary Table 7.** Merqury k-mer (k=21) analysis of Illumina HiSeq reads sequenced from the accession in YUN_Tarv_1.0, showing greater QV scores in T_arvense_v2 for the equivalent top 7 scaffolds based on k-mers found uniquely in each assembly and those shared with the read set.

**Supplementary Table 8.** Merqury k-mer (k=21) analysis of Illumina HiSeq reads (PCR-free) sequenced from the accession MN106-Ref, showing greater QV scores in T_arvense_v2 for the equivalent top 7 scaffolds based on k-mers found uniquely in each assembly and those shared with the read set.

**Supplementary Table 9.** Merqury k-mer (k=21) analysis of each total assembly showing relative completeness of k-mers present in each read set from Illumina HiSeq.

**Supplementary Information 1**. Manual Curation of predicted miRNAs.

**Supplementary File 1.** Normalized read counts for the genes expressed in each of the tissues analyzed (See excel file). Tau values are incorporated in each of the genes to highlight the specificity.

**Supplementary File 2.** Top 30 most expressed genes in each tissue, relative to the mean across all tissues, from the subset of genes with a high/absolute tau specificity score

**Supplementary File 3.** Location of SNPs and the primers used in the genotyping of EMS-based population for development of linkage map.

**Supplementary File 4.** Genetic map developed using an F2 population derived from MN106 and Ames32867.

**Supplementary File 5.** Genetic map developed using an F2 population derived from MN106 and 2019-M2-111.

**Supplementary File 6.** Phenotypes, total reads, and coverage associated with the accessions used for GWAS.

## References

Amselem, J., Cornut, G., Choisne, N., Alaux, M., Alfama-Depauw, F., Jamilloux, V., Maumus, F., Letellier, T., Luyten, I., Pommier, C., Adam-Blondon, A.-F., and Quesneville, H. (2019). RepetDB: a unified resource for transposable element references. Mob. DNA 10: 6.

Andersen, T.G. and Halkier, B.A. (2014). Upon bolting the GTR1 and GTR2 transporters mediate transport of glucosinolates to the inflorescence rather than roots. Plant Signal. Behav. 9: e27740.

Axtell, M.J. (2013a). Classification and comparison of small RNAs from plants. Annu. Rev. Plant Biol. 64: 137–159.

Axtell, M.J. (2013b). ShortStack: comprehensive annotation and quantification of small RNA genes. RNA 19: 740–751.

Bao, W., Kojima, K.K., and Kohany, O. (2015). Repbase Update, a database of repetitive elements in eukaryotic genomes. Mob. DNA 6: 11.

Beilstein, M.A., Nagalingum, N.S., Clements, M.D., Manchester, S.R., and Mathews, S. (2010). Dated molecular phylogenies indicate a Miocene origin for Arabidopsis thaliana. Proc. Natl. Acad. Sci. U. S. A. 107: 18724–18728.

Benson, G. (1999). Tandem repeats finder: a program to analyze DNA sequences. Nucleic Acids Res. 27: 573–580.

Beric, A., Mabry, M.E., Harkess, A.E., Brose, J., Schranz, M.E., Conant, G.C., Edger, P.P., Meyers, B.C., and Pires, J.C. (2021). Comparative phylogenetics of repetitive elements in a diverse order of flowering plants (Brassicales). G3.

Bewick, A.J. et al. (2016). On the origin and evolutionary consequences of gene body DNA methylation. Proc. Natl. Acad. Sci. U. S. A. 113: 9111–9116.

Bewick, A.J. and Schmitz, R.J. (2017). Gene body DNA methylation in plants. Curr. Opin. Plant Biol. 36: 103–110.

Boateng, A.A., Mullen, C.A., and Goldberg, N.M. (2010). Producing Stable Pyrolysis Liquids from the Oil-Seed Presscakes of Mustard Family Plants: Pennycress (Thlaspi arvense L.) and Camelina (Camelina sativa). Energy Fuels 24: 6624–6632.

Boutet, E., Lieberherr, D., Tognolli, M., Schneider, M., and Bairoch, A. (2007). UniProtKB/Swiss-Prot. Methods Mol. Biol. 406: 89–112.

Bradbury, P.J., Zhang, Z., Kroon, D.E., Casstevens, T.M., Ramdoss, Y., and Buckler, E.S. (2007). TASSEL: software for association mapping of complex traits in diverse samples. Bioinformatics 23: 2633–2635.

Bucher, E., Reinders, J., and Mirouze, M. (2012). Epigenetic control of transposon transcription and mobility in Arabidopsis. Curr. Opin. Plant Biol. 15: 503–510.

Campbell, M.S. et al. (2013). MAKER-P: A Tool Kit for the Rapid Creation, Management, and Quality Control of Plant Genome Annotations. Plant Physiol. 164: 513–524.

Campbell, M.S., Holt, C., Moore, B., and Yandell, M. (2014). Genome Annotation and Curation Using MAKER and MAKER-P. Curr. Protoc. Bioinformatics 48: 4.11.1–39.

Catoni, M., Jonesman, T., Cerruti, E., and Paszkowski, J. (2018). Mobilization of Pack-CACTA transposons in Arabidopsis suggests the mechanism of gene shuffling. Nucleic Acids Res. 47: 1311–1320.

Chakraborty, M., Baldwin-Brown, J.G., Long, A.D., and Emerson, J.J. (2016). Contiguous and accurate de novo assembly of metazoan genomes with modest long read coverage. Nucleic Acids Res. 44: e147.

Chopra, R. et al. (2018). Translational genomics using Arabidopsis as a model enables the characterization of pennycress genes through forward and reverse genetics. Plant J. 96: 1093–1105.

Chopra, R., Folstad, N., Lyons, J., and Ulmasov, T. (2019). The adaptable use of Brassica NIRS calibration equations to identify pennycress variants to facilitate the rapid domestication of a new winter oilseed crop. Ind. Crops Prod

Chopra, R., Folstad, N., and Marks, M.D. (2020a). Combined genotype and fatty-acid analysis of single small field pennycress (Thlaspi arvense) seeds increases the throughput for functional genomics and mutant line selection. Ind. Crops Prod. 156: 112823.

Chopra, R., Johnson, E.B., Emenecker, R., and Cahoon, E.B. (2020b). Identification and stacking of crucial traits required for the domestication of pennycress. Nature Food.

Cingolani, P., Platts, A., Wang, L.L., Coon, M., Nguyen, T., Wang, L., Land, S.J., Lu, X., and Ruden, D.M. (2012). A program for annotating and predicting the effects of single nucleotide polymorphisms, SnpEff: SNPs in the genome of Drosophila melanogaster strain w1118; iso-2; iso-3. Fly 6: 80–92.

Claver, A., Rey, R., López, M.V., Picorel, R., and Alfonso, M. (2017). Identification of target genes and processes involved in erucic acid accumulation during seed development in the biodiesel feedstock Pennycress (Thlaspi arvense L.). J. Plant Physiol. 208: 7–16.

Cubins, J.A., Wells, M.S., Frels, K., Ott, M.A., Forcella, F., Johnson, G.A., Walia, M.K., Becker, R.L., and Gesch, R.W. (2019). Management of pennycress as a winter annual cash cover crop. A review. Agron. Sustain. Dev. 39: 46.

Dobin, A., Davis, C.A., Schlesinger, F., Drenkow, J., Zaleski, C., Jha, S., Batut, P., Chaisson, M., and Gingeras, T.R. (2013). STAR: ultrafast universal RNA-seq aligner. Bioinformatics 29: 15–21.

Dorn, K.M., Fankhauser, J.D., Wyse, D.L., and Marks, M.D. (2015). A draft genome of field pennycress (Thlaspi arvense) provides tools for the domestication of a new winter biofuel crop. DNA Res. 22: 121–131.

Du, H. and Liang, C. (2019). Assembly of chromosome-scale contigs by efficiently resolving repetitive sequences with long reads. Nat. Commun. 10: 5360.

Eberle, C.A., Thom, M.D., Nemec, K.T., Forcella, F., Lundgren, J.G., Gesch, R.W., Riedell, W.E., Papiernik, S.K., Wagner, A., Peterson, D.H., and Eklund, J.J. (2015). Using pennycress, camelina, and canola cash cover crops to provision pollinators. Ind. Crops Prod. 75: 20–25.

Esfahanian, M., Nazarenus, T.J., Freund, M.M., McIntosh, G., Phippen, W.B., Phippen, M.E., Durrett, T.P., Cahoon, E.B., and Sedbrook, J.C. (2021). Generating Pennycress (Thlaspi arvense) Seed Triacylglycerols and Acetyl-Triacylglycerols Containing Medium-Chain Fatty Acids. Frontiers in Energy Research 9: 1.

Fan, J., Shonnard, D.R., Kalnes, T.N., Johnsen, P.B., and Rao, S. (2013). A life cycle assessment of pennycress (Thlaspi arvense L.) -derived jet fuel and diesel. Biomass Bioenergy 55: 87– 100.

Flynn, J.M., Hubley, R., Goubert, C., Rosen, J., Clark, A.G., Feschotte, C., and Smit, A.F. (2020). RepeatModeler2 for automated genomic discovery of transposable element families. Proc. Natl. Acad. Sci. U. S. A. 117: 9451–9457.

Franzke, A., Lysak, M.A., Al-Shehbaz, I.A., Koch, M.A., and Mummenhoff, K. (2011). Cabbage family affairs: the evolutionary history of Brassicaceae. Trends Plant Sci. 16: 108–116.

Frels, K., Chopra, R., Dorn, K.M., Wyse, D.L., Marks, M.D., and Anderson, J.A. (2019). Genetic Diversity of Field Pennycress (Thlaspi arvense) Reveals Untapped Variability and Paths Toward Selection for Domestication. Agronomy 9: 302.

Gatter, T. and Stadler, P.F. (2019). Ryūtō: network-flow based transcriptome reconstruction. BMC Bioinformatics 20: 190.

Geng, Y., Guan, Y., Qiong, L., Lu, S., An, M., Crabbe, M.J.C., Qi, J., Zhao, F., Qiao, Q., and Zhang, T. (2021). Genomic analysis of field pennycress (Thlaspi arvense) provides insights into mechanisms of adaptation to high elevation. BMC Biol. 19: 143.

Ghurye, J., Pop, M., Koren, S., Bickhart, D., and Chin, C.-S. (2017). Scaffolding of long read assemblies using long range contact information. BMC Genomics 18: 527.

Guan, D., McCarthy, S.A., Wood, J., Howe, K., Wang, Y., and Durbin, R. (2020). Identifying and removing haplotypic duplication in primary genome assemblies. Bioinformatics 36: 2896– 2898.

Gurevich, A., Saveliev, V., Vyahhi, N., and Tesler, G. (2013). QUAST: quality assessment tool for genome assemblies. Bioinformatics 29: 1072–1075.

Hardcastle, T.J., Müller, S.Y., and Baulcombe, D.C. (2018). Towards annotating the plant epigenome: the Arabidopsis thaliana small RNA locus map. Sci. Rep. 8: 6338.

He, G., Chen, B., Wang, X., Li, X., Li, J., He, H., Yang, M., Lu, L., Qi, Y., Wang, X., and Deng, X.W. (2013). Conservation and divergence of transcriptomic and epigenomic variation in maize hybrids. Genome Biol. 14: R57.

Hill, W.G. and Weir, B.S. (1988). Variances and covariances of squared linkage disequilibria in finite populations. Theor. Popul. Biol. 33: 54–78.

Jarvis, B.A., Romsdahl, T.B., McGinn, M.G., Nazarenus, T.J., Cahoon, E.B., Chapman, K.D., and Sedbrook, J.C. (2021). CRISPR/Cas9-Induced fad2 and rod1 Mutations Stacked With fae1 Confer High Oleic Acid Seed Oil in Pennycress (Thlaspi arvense L.). Front. Plant Sci. 12: 652319.

Johnson, G.A., Kantar, M.B., Betts, K.J., and Wyse, D.L. (2015). Field Pennycress Production and Weed Control in a Double Crop System with Soybean in Minnesota. Agron. J. 107: 532– 540.

Jones, P. et al. (2014). InterProScan 5: genome-scale protein function classification. Bioinformatics 30: 1236–1240.

Kamvar, Z.N., Tabima, J.F., and Grünwald, N.J. (2014). Poppr: an R package for genetic analysis of populations with clonal, partially clonal, and/or sexual reproduction. PeerJ 2: e281.

Koren, S., Walenz, B.P., Berlin, K., Miller, J.R., Bergman, N.H., and Phillippy, A.M. (2017). Canu: scalable and accurate long-read assembly via adaptive k-mer weighting and repeat separation. Genome Res. 27: 722–736.

Korf, I. (2004). Gene finding in novel genomes. BMC Bioinformatics 5: 59.

Lagesen, K., Hallin, P., Rødland, E.A., Staerfeldt, H.-H., Rognes, T., and Ussery, D.W. (2007). RNAmmer: consistent and rapid annotation of ribosomal RNA genes. Nucleic Acids Res. 35: 3100–3108.

Lam, E.T., Hastie, A., Lin, C., Ehrlich, D., Das, S.K., Austin, M.D., Deshpande, P., Cao, H., Nagarajan, N., Xiao, M., and Kwok, P.-Y. (2012). Genome mapping on nanochannel arrays for structural variation analysis and sequence assembly. Nat. Biotechnol. 30: 771–776.

Liao, Y., Smyth, G.K., and Shi, W. (2014). featureCounts: an efficient general purpose program for assigning sequence reads to genomic features. Bioinformatics 30: 923–930.

Li, H. (2018). Minimap2: pairwise alignment for nucleotide sequences. Bioinformatics 34: 3094–3100.

Li, H. and Durbin, R. (2009). Fast and accurate short read alignment with Burrows–Wheeler transform. Bioinformatics 25: 1754–1760.

Li, H., Handsaker, B., Wysoker, A., Fennell, T., Ruan, J., Homer, N., Marth, G., Abecasis, G., Durbin, R., and 1000 Genome Project Data Processing Subgroup (2009). The Sequence Alignment/Map format and SAMtools. Bioinformatics 25: 2078–2079.

Liu, X., Huang, M., Fan, B., Buckler, E.S., and Zhang, Z. (2016). Iterative Usage of Fixed and Random Effect Models for Powerful and Efficient Genome-Wide Association Studies. PLoS Genet. 12: e1005767.

Lowe, T.M. and Eddy, S.R. (1997). tRNAscan-SE: a program for improved detection of transfer RNA genes in genomic sequence. Nucleic Acids Res. 25: 955–964.

Lu, K. et al. (2019). Whole-genome resequencing reveals Brassica napus origin and genetic loci involved in its improvement. Nat. Commun. 10: 1154.

Lunardon, A., Johnson, N.R., Hagerott, E., Phifer, T., Polydore, S., Coruh, C., and Axtell, M.J. (2020). Integrated annotations and analyses of small RNA–producing loci from 47 diverse plants. Genome Res. 30: 497–513.

Marçais, G. and Kingsford, C. (2011). A fast, lock-free approach for efficient parallel counting of occurrences of k-mers. Bioinformatics 27: 764–770.

Marks, M.D., Chopra, R., and Sedbrook, J.C. (2021). Technologies enabling rapid crop improvements for sustainable agriculture: example pennycress (Thlaspi arvense L.). Emerg Top Life Sci 5: 325–335.

Martin, M. (2011). Cutadapt removes adapter sequences from high-throughput sequencing reads. EMBnet.journal 17: 10–12.

McGinn, M. et al. (2019). Molecular tools enabling pennycress (Thlaspi arvense) as a model plant and oilseed cash cover crop. Plant Biotechnol. J. 17: 776–788.

Meinke, D.W., Cherry, J.M., Dean, C., Rounsley, S.D., and Koornneef, M. (1998). Arabidopsis thaliana: a model plant for genome analysis. Science 282: 662, 679–82.

Michalovova, M., Vyskot, B., and Kejnovsky, E. (2013). Analysis of plastid and mitochondrial DNA insertions in the nucleus (NUPTs and NUMTs) of six plant species: size, relative age and chromosomal localization. Heredity 111: 314–320.

Michelmore, R.W., Paran, I., and Kesseli, R.V. (1991). Identification of markers linked to disease-resistance genes by bulked segregant analysis: a rapid method to detect markers in specific genomic regions by using segregating populations. Proc. Natl. Acad. Sci. U. S. A. 88: 9828–9832.

Moore, S.A., Wells, M.S., Gesch, R.W., Becker, R.L., Rosen, C.J., and Wilson, M.L. (2020). Pennycress as a Cash Cover-Crop: Improving the Sustainability of Sweet Corn Production Systems. Agronomy 10: 614.

Moser, B.R. (2012). Biodiesel from alternative oilseed feedstocks: camelina and field pennycress. Biofuels 3: 193–209.

Moser, B.R., Knothe, G., Vaughn, S.F., and Isbell, T.A. (2009). Production and Evaluation of Biodiesel from Field Pennycress (Thlaspi arvense L.) Oil. Energy Fuels 23: 4149–4155.

Mulligan, G.A. (1957). CHROMOSOME NUMBERS OF CANADIAN WEEDS. I. Can. J. Bot. 35: 779– 789.

Mulligan, G.A. and Kevan, P.G. (1973). Color, brightness, and other floral characteristics attracting insects to the blossoms of some Canadian weeds. Can. J. Bot. 51: 1939–1952.

Murat, F., Louis, A., Maumus, F., Armero, A., Cooke, R., Quesneville, H., Roest Crollius, H., and Salse, J. (2015). Understanding Brassicaceae evolution through ancestral genome reconstruction. Genome Biol. 16: 262.

Nawrocki, E.P. and Eddy, S.R. (2013). Infernal 1.1: 100-fold faster RNA homology searches. Bioinformatics 29: 2933–2935.

Niederhuth, C.E. et al. (2016). Widespread natural variation of DNA methylation within angiosperms. Genome Biol. 17: 194.

Niittylä, T., Messerli, G., Trevisan, M., Chen, J., Smith, A.M., and Zeeman, S.C. (2004). A Previously Unknown Maltose Transporter Essential for Starch Degradation in Leaves. Science 303: 87–89.

Niknafs, Y.S., Pandian, B., Iyer, H.K., Chinnaiyan, A.M., and Iyer, M.K. (2017). TACO produces robust multisample transcriptome assemblies from RNA-seq. Nat. Methods 14: 68–70.

Nour-Eldin, H.H., Andersen, T.G., Burow, M., Madsen, S.R., Jorgensen, M.E., Olsen, C.E., and Dreyer, I. (2012). NRT/PTR transporters are essential for translocation of glucosinolate defence compounds to seeds. Nature 488: 531+.

Ott, M.A., Eberle, C.A., Thom, M.D., Archer, D.W., Forcella, F., Gesch, R.W., and Wyse, D.L. (2019). Economics and agronomics of relay-cropping pennycress and Camelina with soybean in Minnesota. Agron. J. 111: 1281–1292.

Ou, S. et al. (2019). Benchmarking transposable element annotation methods for creation of a streamlined, comprehensive pipeline. Genome Biol. 20: 275.

Patra Bhattacharya, D., Canzler, S., Kehr, S., Hertel, J., Grosse, I., and Stadler, P.F. (2016). Phylogenetic distribution of plant snoRNA families. BMC Genomics 17: 969.

Pedersen, B.S., Eyring, K., De, S., Yang, I.V., and Schwartz, D.A. (2014). Fast and accurate alignment of long bisulfite-seq reads. arXiv [q-bio.GN].

Phippen, W.B. and Phippen, M.E. (2012). Soybean Seed Yield and Quality as a Response to Field Pennycress Residue. Crop Sci. 52: 2767–2773.

Pritchard, J.K., Stephens, M., and Donnelly, P. (2000). Inference of Population Structure Using Multilocus Genotype Data. Genetics 155: 945–959.

Pruitt, K.D., Tatusova, T., Brown, G.R., and Maglott, D.R. (2012). NCBI Reference Sequences (RefSeq): current status, new features and genome annotation policy. Nucleic Acids Res. 40: D130–5.

Purcell, S., Neale, B., Todd-Brown, K., Thomas, L., Ferreira, M.A.R., Bender, D., Maller, J., Sklar, P., de Bakker, P.I.W., Daly, M.J., and Sham, P.C. (2007). PLINK: a tool set for whole-genome association and population-based linkage analyses. Am. J. Hum. Genet. 81: 559– 575.

Quinlan, A.R. and Hall, I.M. (2010). BEDTools: a flexible suite of utilities for comparing genomic features. Bioinformatics 26: 841–842.

Ren, S., Bertels, K., and Al-Ars, Z. (2018). Efficient Acceleration of the Pair-HMMs Forward Algorithm for GATK HaplotypeCaller on Graphics Processing Units. Evol. Bioinform. Online 14: 1176934318760543.

Rhie, A., Walenz, B.P., Koren, S., and Phillippy, A.M. (2020). Merqury: reference-free quality, completeness, and phasing assessment for genome assemblies. Genome Biol. 21: 245.

Robinson, M.D., McCarthy, D.J., and Smyth, G.K. (2010). edgeR: a Bioconductor package for differential expression analysis of digital gene expression data. Bioinformatics 26: 139– 140.

Robinson, M.D. and Oshlack, A. (2010). A scaling normalization method for differential expression analysis of RNA-seq data. Genome Biol. 11: R25.

Scheben, A., Yuan, Y., and Edwards, D. (2016). Advances in genomics for adapting crops to climate change. Current Plant Biology 6: 2–10.

Schiessl, S.V., Quezada-Martinez, D., Tebartz, E., Snowdon, R.J., and Qian, L. (2019). The vernalisation regulator FLOWERING LOCUS C is differentially expressed in biennial and annual Brassica napus. Sci. Rep. 9: 1–15.

Schranz, M.E., Lysak, M.A., and Mitchell-Olds, T. (2006). The ABC’s of comparative genomics in the Brassicaceae: building blocks of crucifer genomes. Trends Plant Sci. 11: 535–542.

Sedbrook, J.C., Phippen, W.B., and Marks, M.D. (2014). New approaches to facilitate rapid domestication of a wild plant to an oilseed crop: example pennycress (Thlaspi arvense L.). Plant Sci. 227: 122–132.

Shared-Protocol-Guidelines-for-Using-a-Salt-Chloroform-Wash-to-Clean-Up-gDNA.pdf

Shumate, A. and Salzberg, S.L. (2020). Liftoff: accurate mapping of gene annotations. Bioinformatics.

Sigman, M.J. and Slotkin, R.K. (2016). The First Rule of Plant Transposable Element Silencing: Location, Location, Location. Plant Cell 28: 304–313.

Simão, F.A., Waterhouse, R.M., Ioannidis, P., Kriventseva, E.V., and Zdobnov, E.M. (2015). BUSCO: assessing genome assembly and annotation completeness with single-copy orthologs. Bioinformatics 31: 3210–3212.

Smit, A.F.A. (2004). Repeat-Masker Open-3.0. http://www.repeatmasker.org.

Smit, A.F.A. and Hubley, R. (2008). RepeatModeler Open-1.0.

Sonnhammer, E.L. and Durbin, R. (1995). A dot-matrix program with dynamic threshold control suited for genomic DNA and protein sequence analysis. Gene 167: GC1–10.

Stam, P. (1993). Construction of integrated genetic linkage maps by means of a new computer package: Join Map. Plant J. 3: 739–744.

Stanke, M., Keller, O., Gunduz, I., Hayes, A., Waack, S., and Morgenstern, B. (2006). AUGUSTUS: ab initio prediction of alternative transcripts. Nucleic Acids Res. 34: W435–9.

Sugihara, Y., Young, L., Yaegashi, H., Natsume, S., Shea, D.J., Takagi, H., Booker, H., Innan, H., Terauchi, R., and Abe, A. (2020). High-performance pipeline for MutMap and QTL-seq. bioRxiv: 2020.06.28.176586.

Sultana, T., Zamborlini, A., Cristofari, G., and Lesage, P. (2017). Integration site selection by retroviruses and transposable elements in eukaryotes. Nat. Rev. Genet. 18: 292–308.

Tadege, M., Sheldon, C.C., Helliwell, C.A., Stoutjesdijk, P., Dennis, E.S., and Peacock, W.J. (2001). Control of flowering time by FLC orthologues in Brassica napus. Plant J. 28: 545– 553.

Tang, H., Zhang, X., Miao, C., Zhang, J., Ming, R., Schnable, J.C., Schnable, P.S., Lyons, E., and Lu, J. (2015). ALLMAPS: robust scaffold ordering based on multiple maps. Genome Biol. 16: 3.

Tardaguila, M. et al. (2018). SQANTI: extensive characterization of long-read transcript sequences for quality control in full-length transcriptome identification and quantification. Genome Res.

Thomas, J.B., Hampton, M.E., Dorn, K.M., David Marks, M., and Carter, C.J. (2017). The pennycress (Thlaspi arvense L.) nectary: structural and transcriptomic characterization. BMC Plant Biol. 17: 201.

Tomato Genome Consortium (2012). The tomato genome sequence provides insights into fleshy fruit evolution. Nature 485: 635–641.

Vaser, R., Sović, I., Nagarajan, N., and Šikić, M. (2017). Fast and accurate de novo genome assembly from long uncorrected reads. Genome Res. 27: 737–746.

Voinnet, O. (2009). Origin, biogenesis, and activity of plant microRNAs. Cell 136: 669–687.

Walker, B.J., Abeel, T., Shea, T., Priest, M., Abouelliel, A., Sakthikumar, S., Cuomo, C.A., Zeng, Q., Wortman, J., Young, S.K., and Earl, A.M. (2014). Pilon: an integrated tool for comprehensive microbial variant detection and genome assembly improvement. PLoS One 9: e112963.

Wang, X. et al. (2011). The genome of the mesopolyploid crop species Brassica rapa. Nat. Genet. 43: 1035–1039.

Warwick, S.I., Francis, A., and Susko, D.J. (2002). The biology of Canadian weeds. 9. Thlaspi arvense L. (updated). Can. J. Plant Sci. 82: 803–823.

Weyers, S.L., Gesch, R.W., Forcella, F., Eberle, C.A., Thom, M.D., Matthees, H.L., Ott, M., Feyereisen, G.W., and Strock, J.S. (2021). Surface runoff and nutrient dynamics in cover crop-soybean systems in the Upper Midwest. J. Environ. Qual. 50: 158–171.

Weyers, S., Thom, M., Forcella, F., Eberle, C., Matthees, H., Gesch, R., Ott, M., Feyereisen, G., Strock, J., and Wyse, D. (2019). Reduced Potential for Nitrogen Loss in Cover Crop-Soybean Relay Systems in a Cold Climate. J. Environ. Qual. 48: 660–669.

Workman, R., Fedak, R., Kilburn, D., Hao, S., Liu, K., and Timp, W. (2019). High molecular weight DNA extraction from recalcitrant plant species for third generation sequencing v1 (protocols.Io.4vbgw2n). protocols.io.

Wu, D. et al. (2019). Whole-Genome Resequencing of a Worldwide Collection of Rapeseed Accessions Reveals the Genetic Basis of Ecotype Divergence. Mol. Plant 12: 30–43.

Yanai, I., Benjamin, H., Shmoish, M., Chalifa-Caspi, V., Shklar, M., Ophir, R., Bar-Even, A., Horn-Saban, S., Safran, M., Domany, E., Lancet, D., and Shmueli, O. (2005). Genome-wide midrange transcription profiles reveal expression level relationships in human tissue specification. Bioinformatics 21: 650–659.

Yang, R. et al. (2013). The Reference Genome of the Halophytic Plant Eutrema salsugineum. Front. Plant Sci. 4: 46.

Yin, L., Zhang, H., Tang, Z., Xu, J., Yin, D., Zhang, Z., Yuan, X., Zhu, M., Zhao, S., Li, X., and Liu, X. (2021). rMVP: A Memory-efficient, Visualization-enhanced, and Parallel-accelerated tool for Genome-Wide Association Study. Genomics Proteomics Bioinformatics.

Zhang, H., Lang, Z., and Zhu, J.-K. (2018). Dynamics and function of DNA methylation in plants. Nat. Rev. Mol. Cell Biol. 19: 489–506.

Zhang, S.-J., Liu, L., Yang, R., and Wang, X. (2020). Genome Size Evolution Mediated by Gypsy Retrotransposons in Brassicaceae. Genomics Proteomics Bioinformatics 18: 321–332.

Zhang, X., Yazaki, J., Sundaresan, A., Cokus, S., Chan, S.W.-L., Chen, H., Henderson, I.R., Shinn, P., Pellegrini, M., Jacobsen, S.E., and Ecker, J.R. (2006). Genome-wide high-resolution mapping and functional analysis of DNA methylation in arabidopsis. Cell 126: 1189–1201.

Zilberman, D., Gehring, M., Tran, R.K., Ballinger, T., and Henikoff, S. (2006). Genome-wide analysis of Arabidopsis thaliana DNA methylation uncovers an interdependence between methylation and transcription. Nat. Genet. 39: 61–69.

Zou, C., Lehti-Shiu, M.D., Thibaud-Nissen, F., Prakash, T., Buell, C.R., and Shiu, S.-H. (2009). Evolutionary and expression signatures of pseudogenes in Arabidopsis and rice. Plant Physiol. 151: 3–15.

