## Supplemental Figures and Tables for "Chromosome-level *Thlaspi arvense* genome provides new tools for translational research and for a newly domesticated cash cover crop of the cooler climates"

### Content:

### Supplemental Figures 1-16

### Supplemental Tables 1-9

### Supplemental Methods

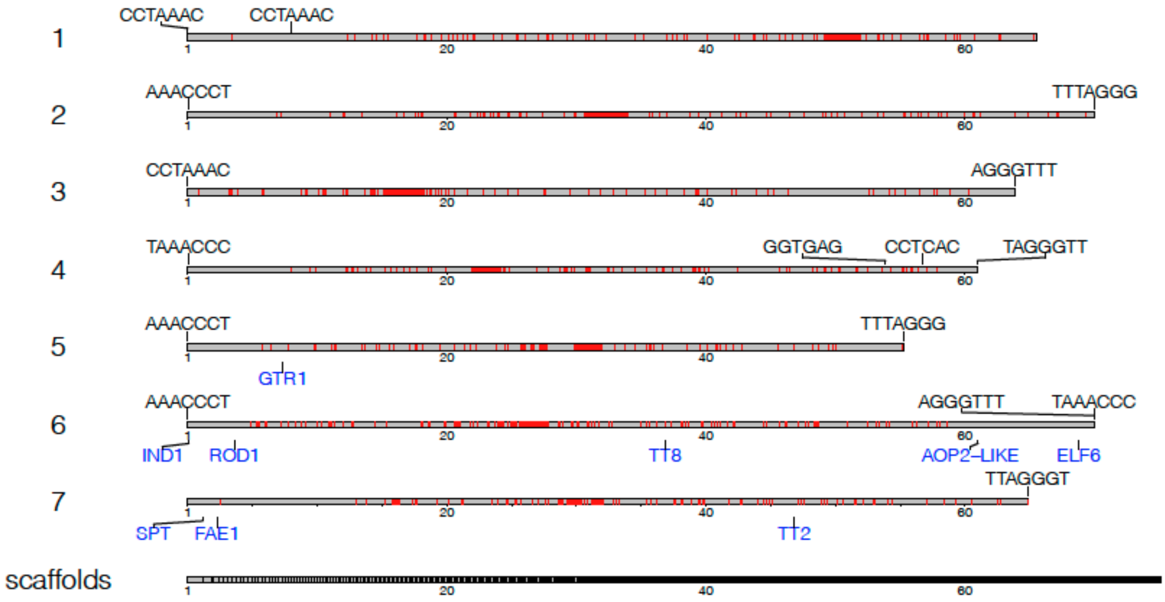


**Supplementary Figure 1.** Karyotype plot of the seven largest scaffolds representing chromosomes in T. arvense MN106-Ref (T_arvense_v2), alongside a concatenation of all minor scaffolds. Transposable element LTR annotations (red) are highlighted as an approximate localisation of each centromere, alongside loci containing putative telomeric repeat motifs (black labels) and genes of interest (blue labels) in the *de novo* domestication of pennycress. Scaling is given in Mbp.

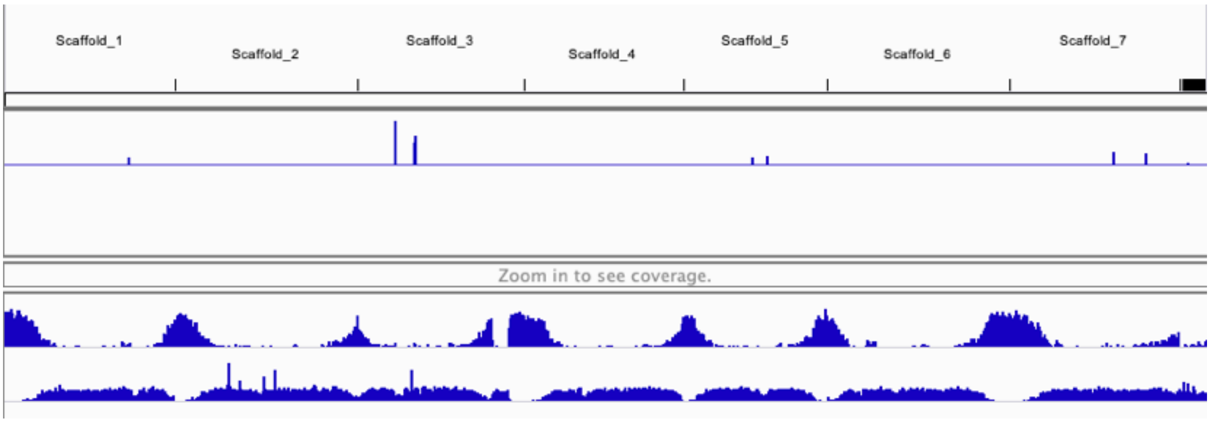


**Supplementary Figure 2.** Integrative Genome Viewer (IGV) snapshot of PacBio read coverage (top track) over the largest seven scaffolds of the genome, including distributions of genes (middle track) and transposable elements (bottom track). Spikes in coverage in scaffolds 1, 3, 5, and 7 are indicative of collapsed repeats, which are typically larger than the average read length.

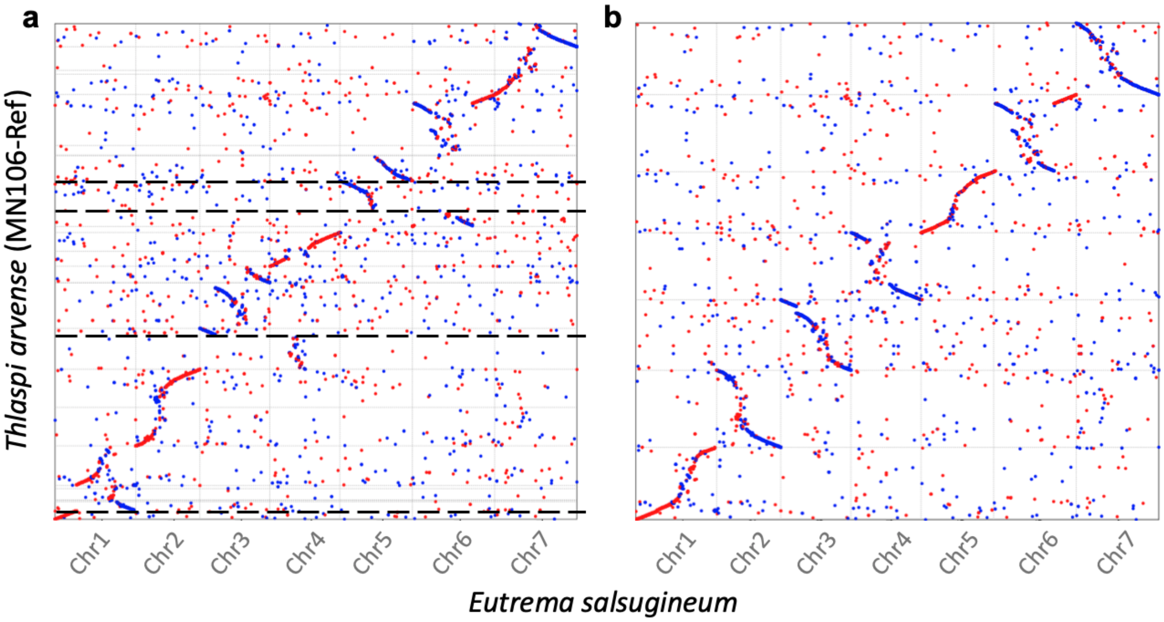


**Supplementary Figure 3.** Sequence dot plots showing the largest seven scaffolds of the closely-related species *E. salsugineum* and their equivalent in *T. arvense* var. MN106-Ref (T_arvense_v2), comparing the difference both **a)** before and **b)** after re-scaffolding. Horizontal dashed lines in (a) denote breakpoints which were manually introduced to the genome based on evaluation of genetic maps, synteny maps and Hi-C data.

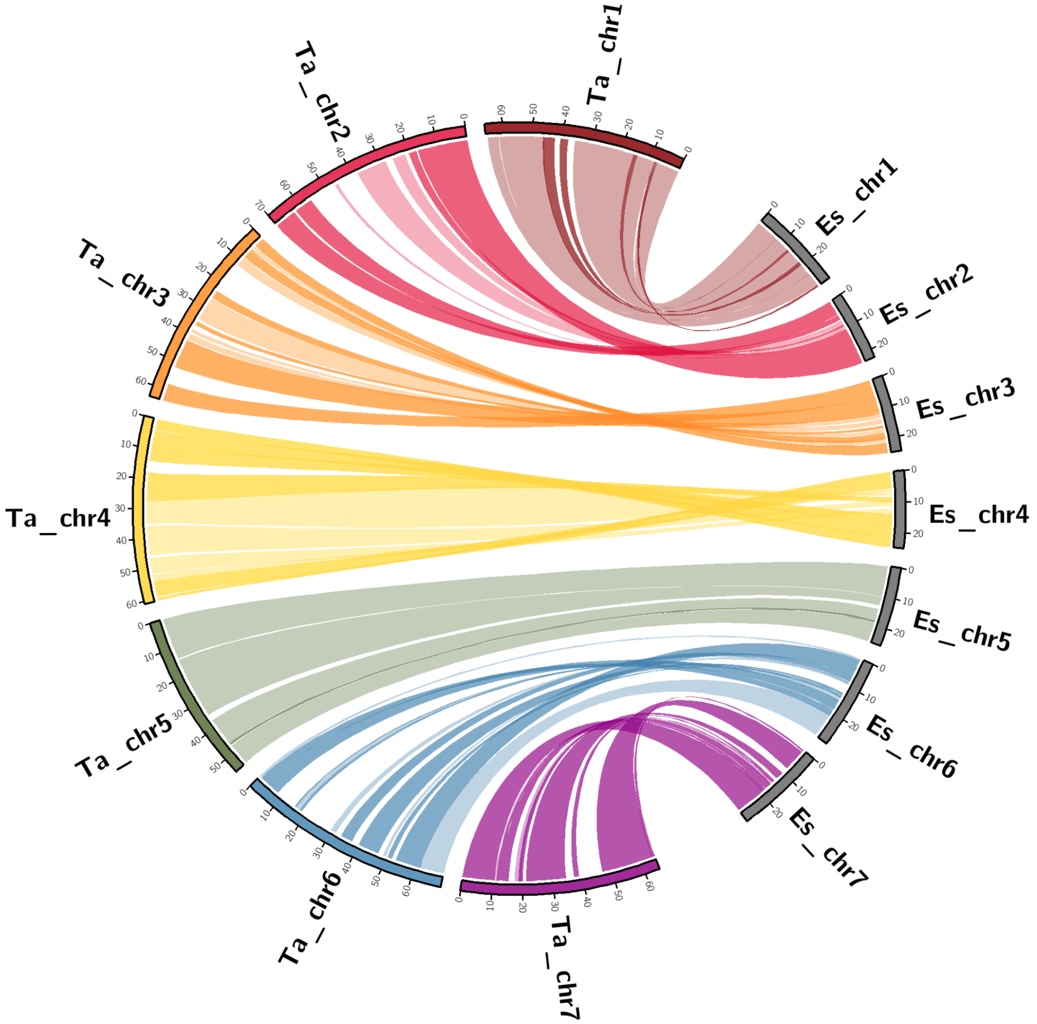


**Supplementary Figure 4.** Synteny analysis between the largest seven scaffolds of the closely-related species *E. salsugineum* and their equivalent in *T. arvense* var. MN106-Ref (T_arvense_v2). Ribbons show the syntenic relationships between the two genomes. Dark ribbons indicate syntenic blocks in inverse orientation.

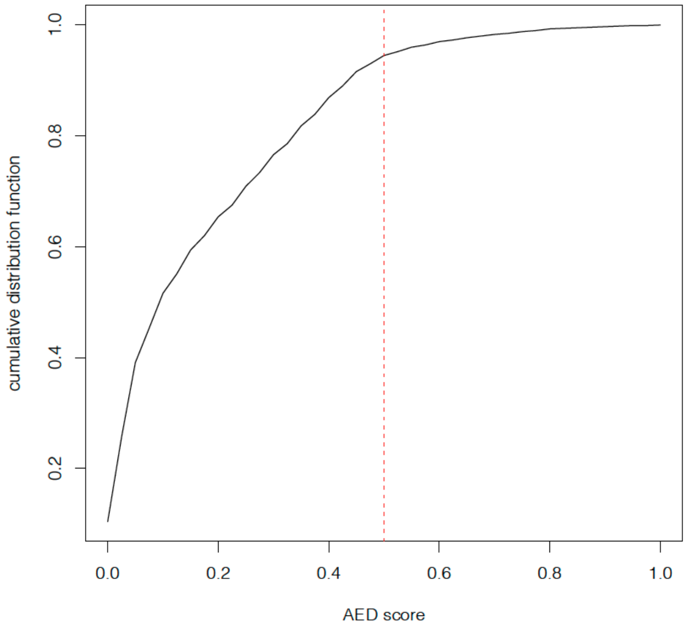


**Supplementary Figure 5.** The cumulative distribution of annotation edit distance (AED) scores from the final set of protein-coding loci, denoting that ~95% of annotated genes are supported with a score ≤ 0.5 overall.

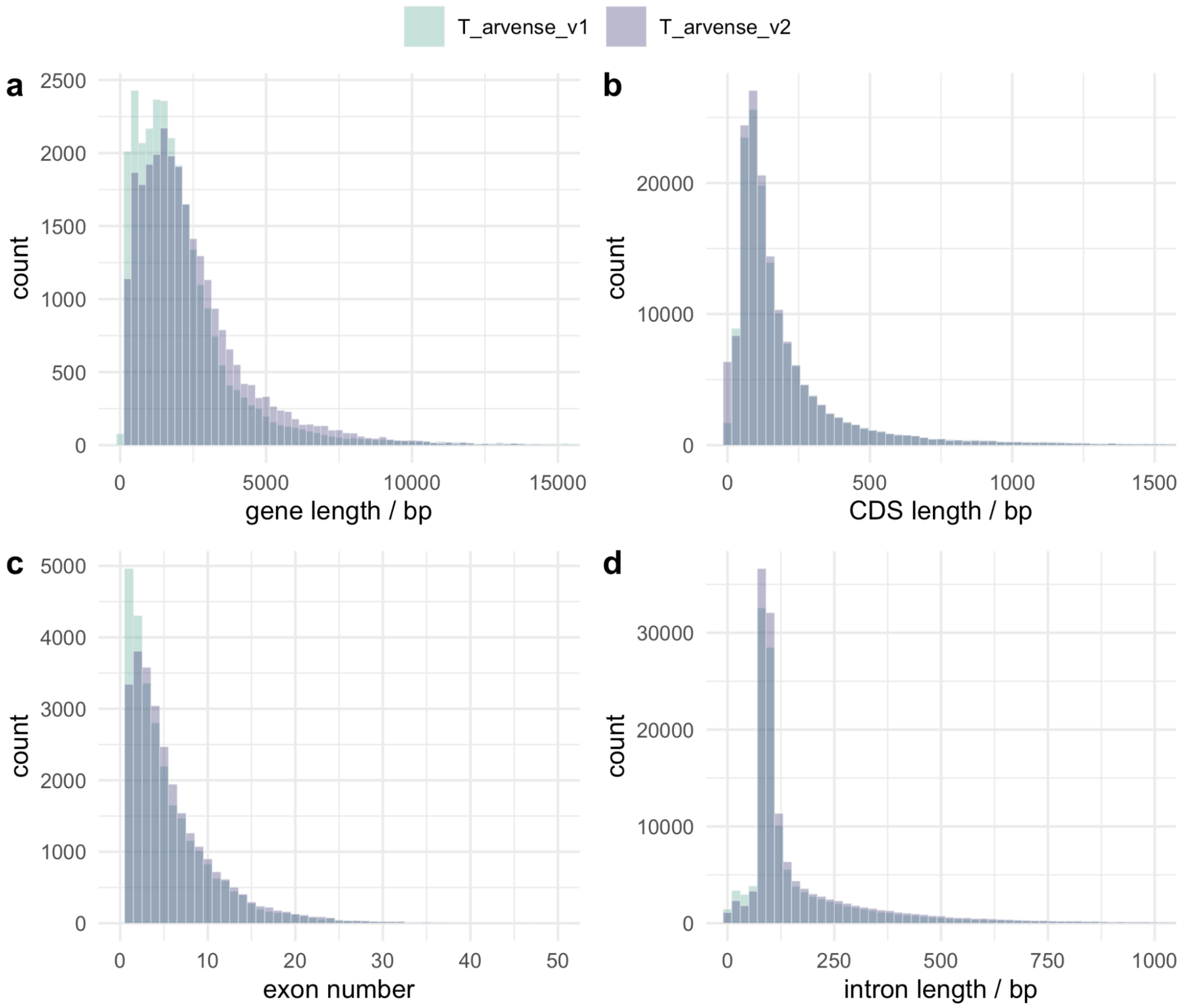


**Supplementary Figure 6.** An overview of annotated genomic feature distributions in comparison to T_arvense_v1 for **a)** gene lengths, **b)** CDS lengths, **c)** per gene exon number, and **d)** intron lengths.

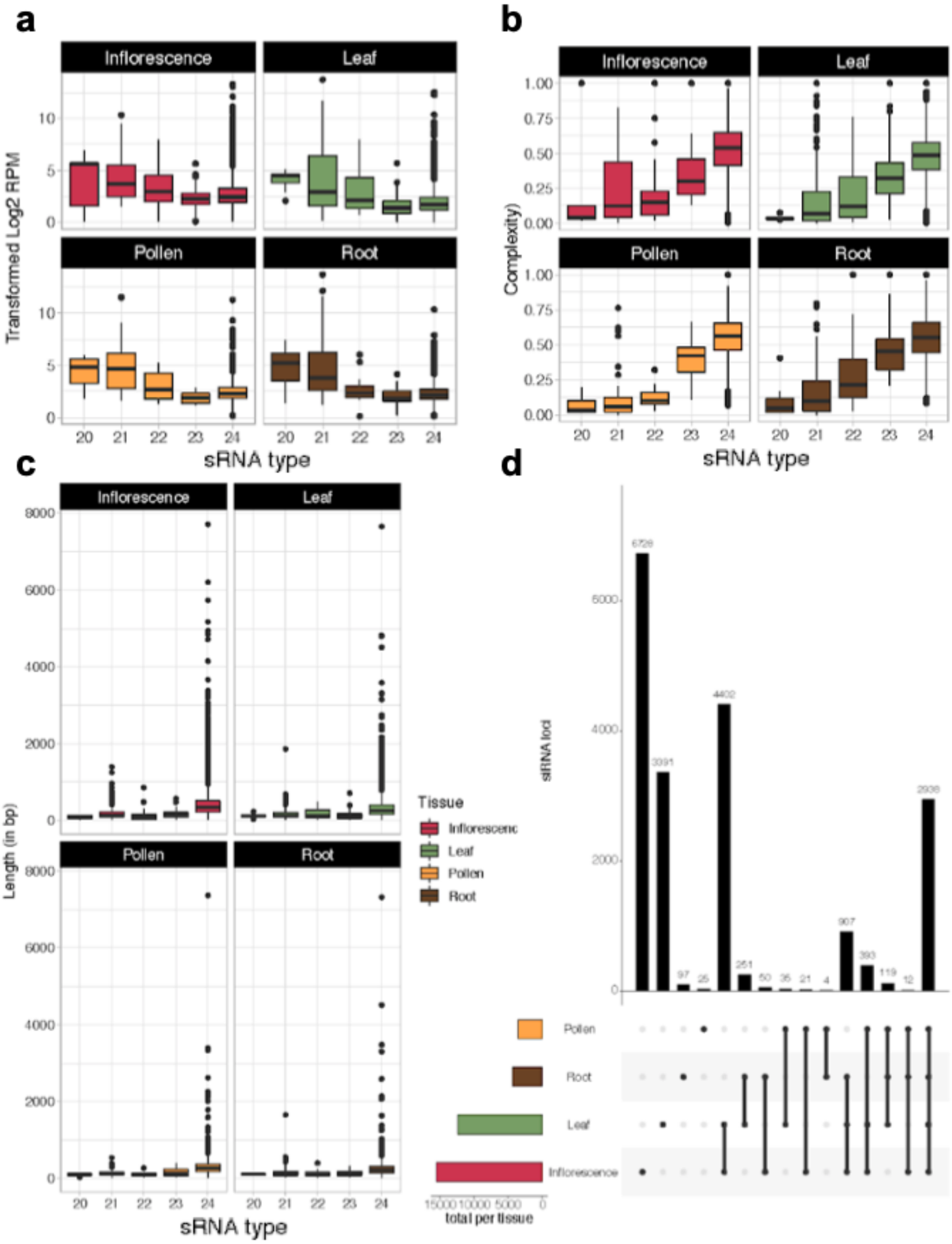


**Supplementary Figure 7.** Small RNA (sRNA) annotation in the T_arvense_v2 genome assembly. **a)** sRNA loci per tissue of origin (RPM = reads per million). **b)** sRNA complexity, measured as "number of distinct alignments / total number of alignments". **c)** sRNA locus size distribution. **d)** Co-occurrence of sRNAs between tissues. Colored horizontal bars show the total number of loci per tissue.

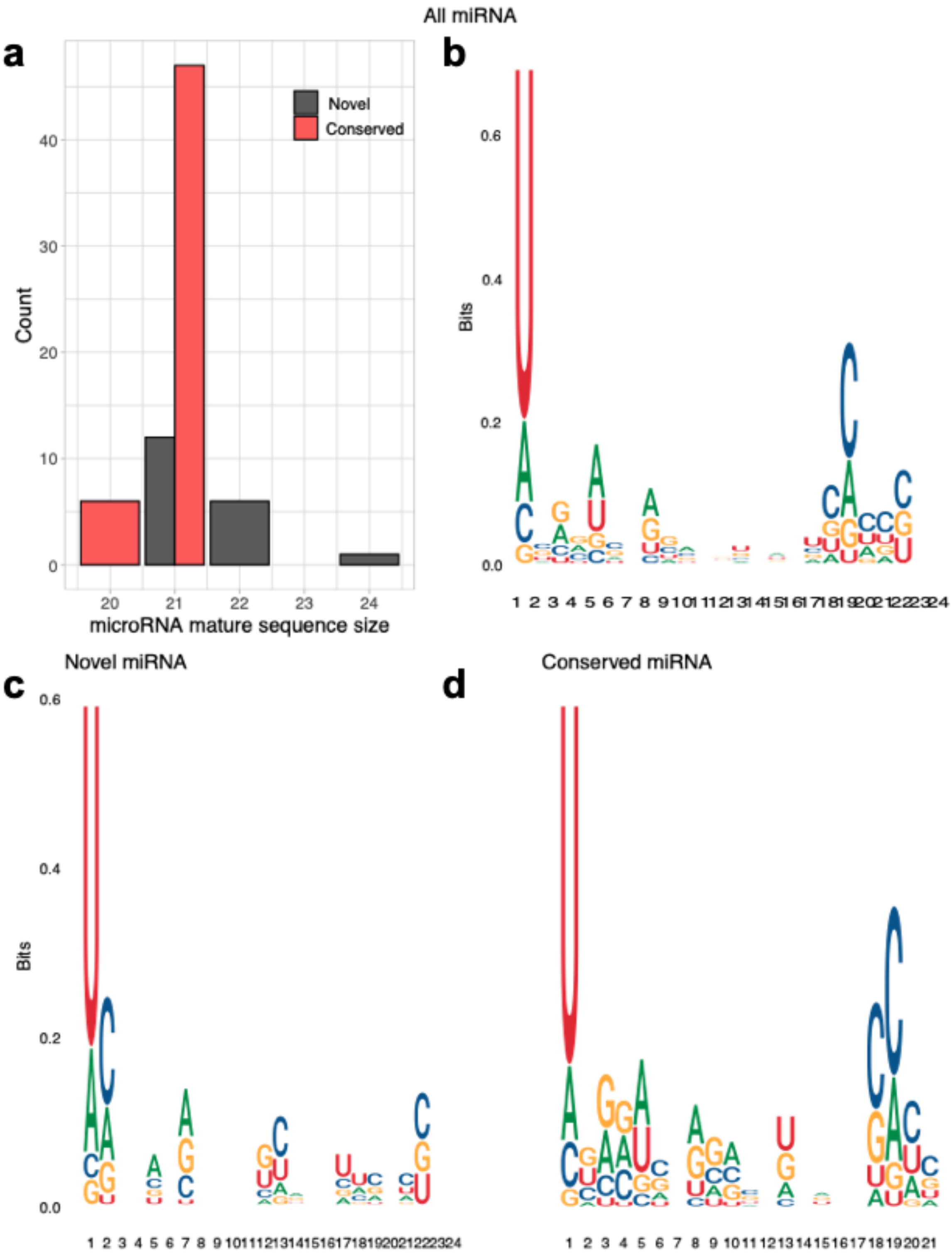


**Supplementary Figure 8.** MiRNAs in the T_arvense_v2 genome assembly. **a)** Size of mature miRNAs identified in this study, split into novel and conserved miRNA species. **b-d)** Sequence conservation in miRNAs, measured in bits foreach position of the mature microRNA (Schneider and Stephens 1990). Sequences were aligned from the 5'-end. Different panels show sequence conservation of all miRNAs **(b)**, only novel miRNAs **(c)** and only conserved miRNAs **(d)**.

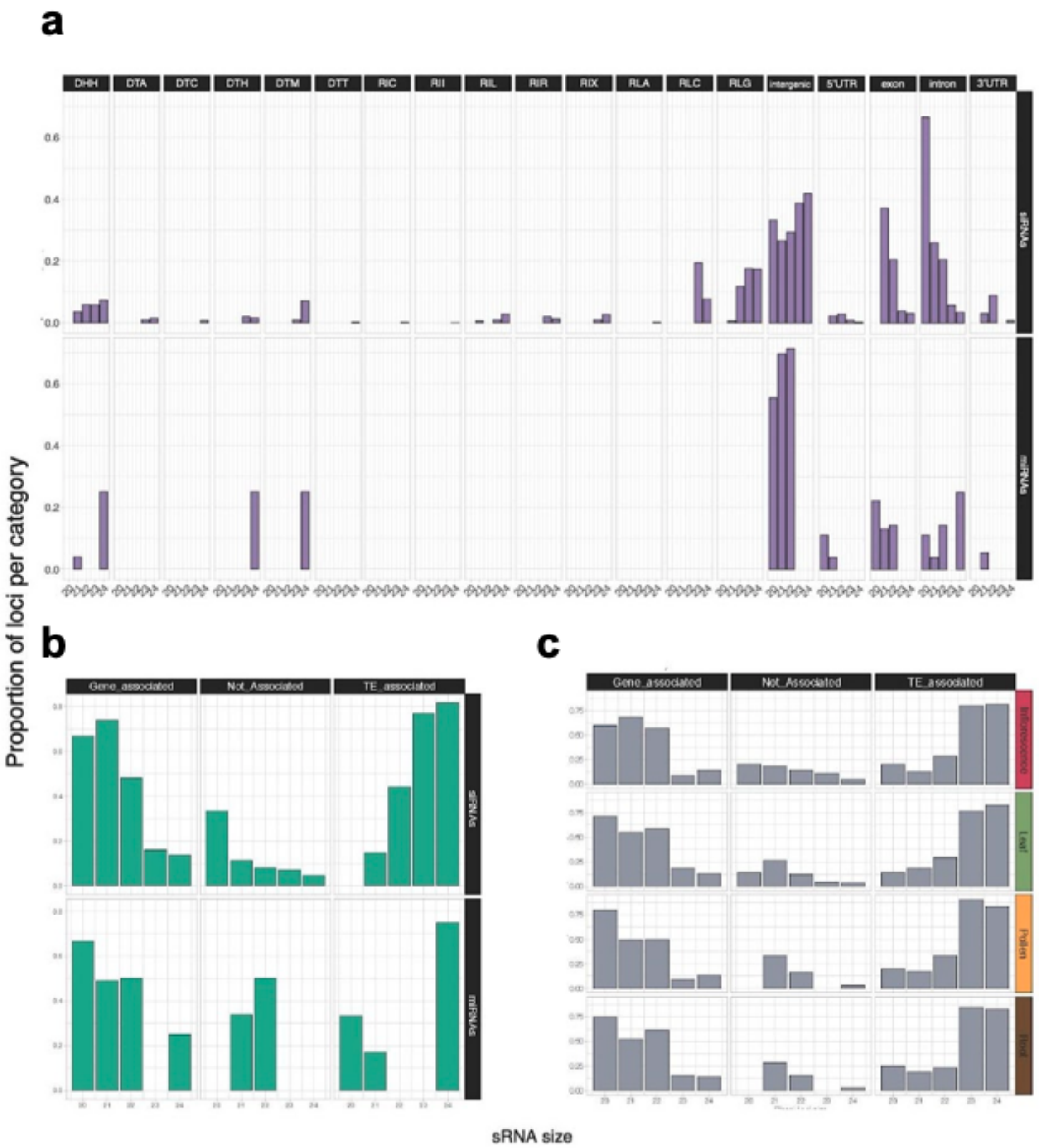


**Supplementary Figure 9.** sRNA types and their association with different genomic features. **a)** Occupancy of all annotated siRNAs and miRNAs in either genes, TE superfamilies, or intergenic regions. **b,c)** Association of sRNA loci with either TEs or genes within 1.5 Kb distance, for all sRNAs **(b)** and for only phased loci **(c)**.

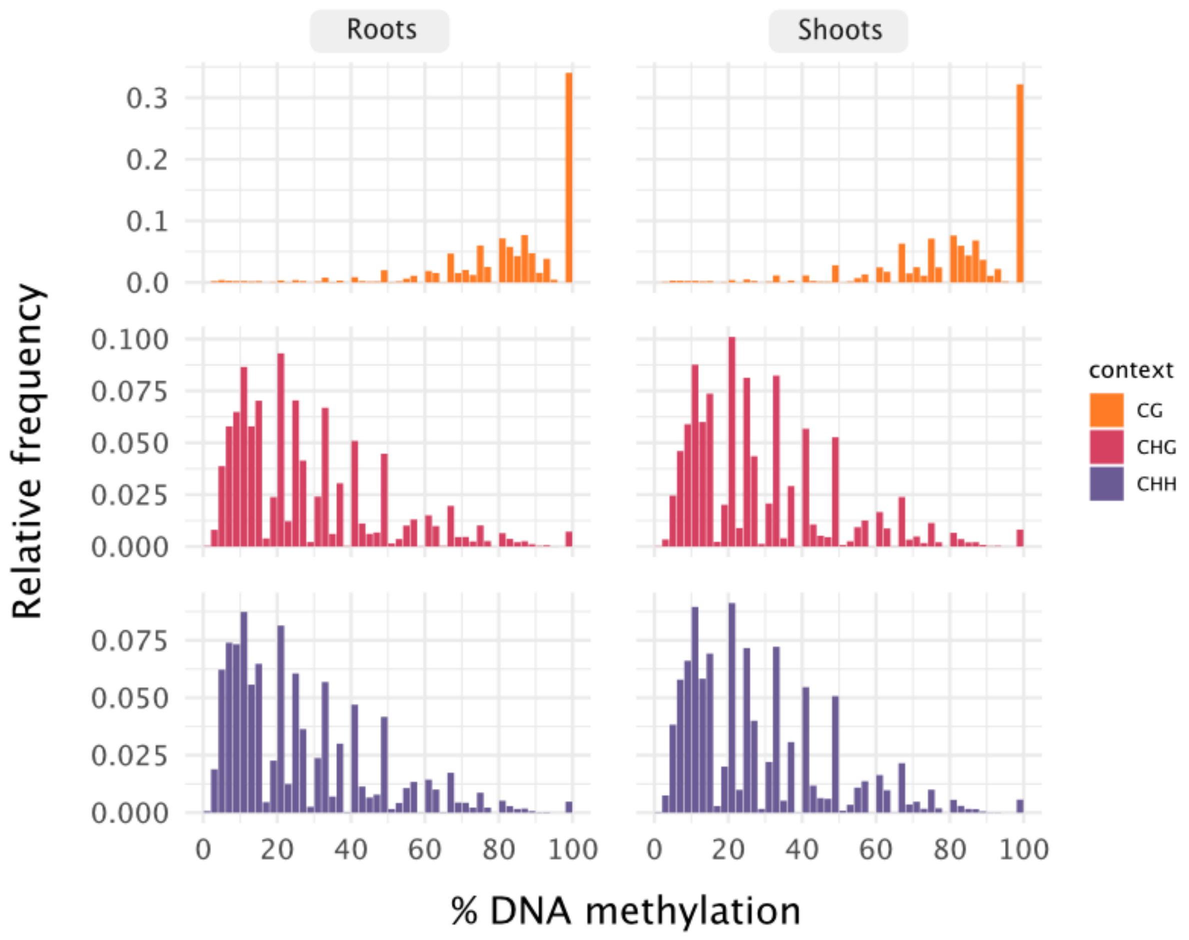


**Supplementary Figure 10.** Methylation rate frequency distribution by sequence context in shoot and root tissues.

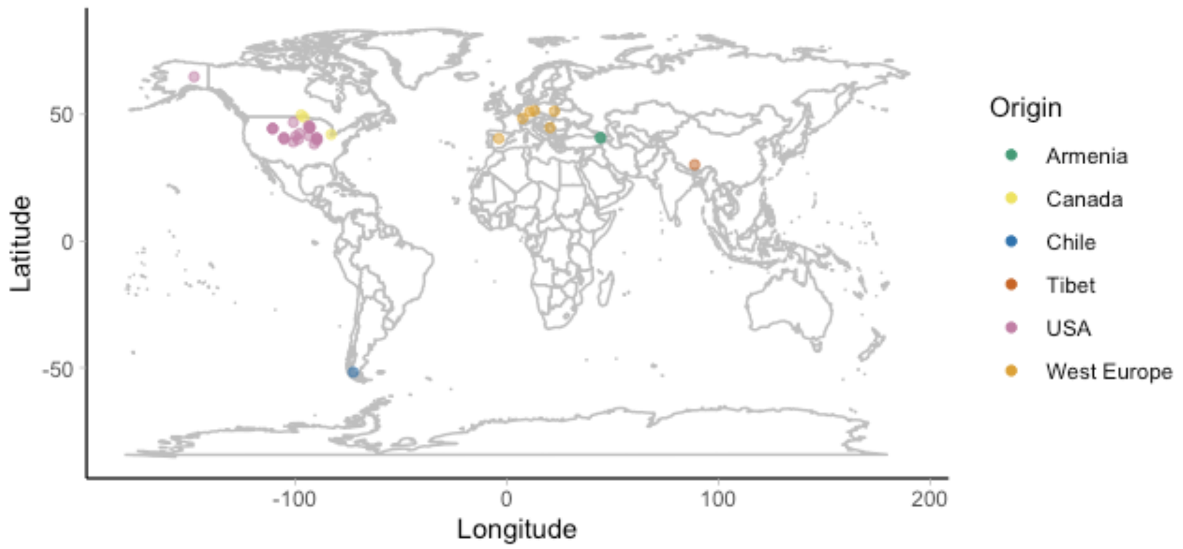


**Supplementary Figure 11.** Map showing original sampling sites of pennycress accessions used for resequencing analysis in this study.

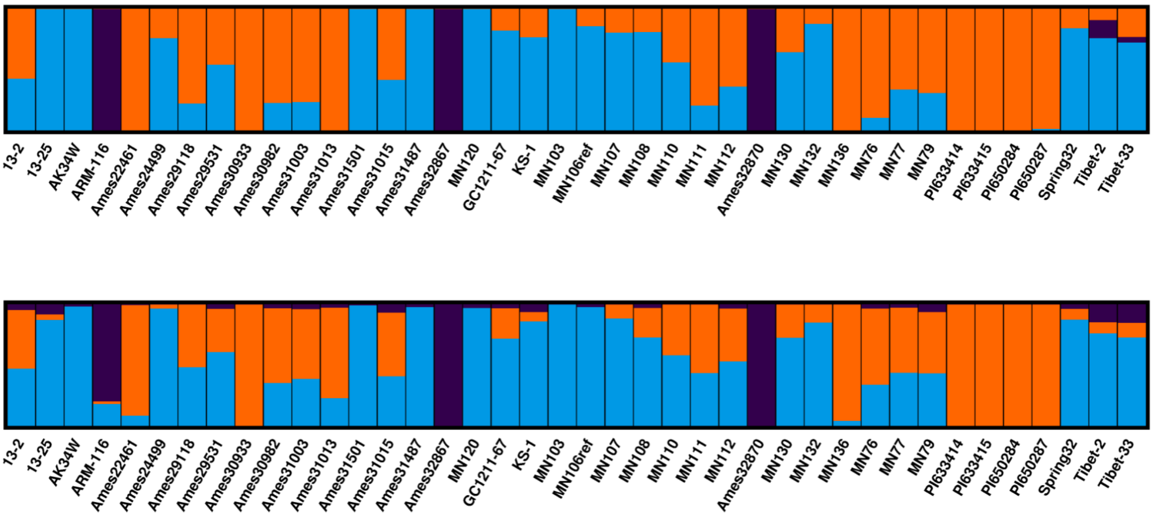


**Supplementary Figure 12.** Structure plot showing inferred population membership for SNP data (top) and Indel data (bottom) at k = 3 for the re-sequenced accessions.

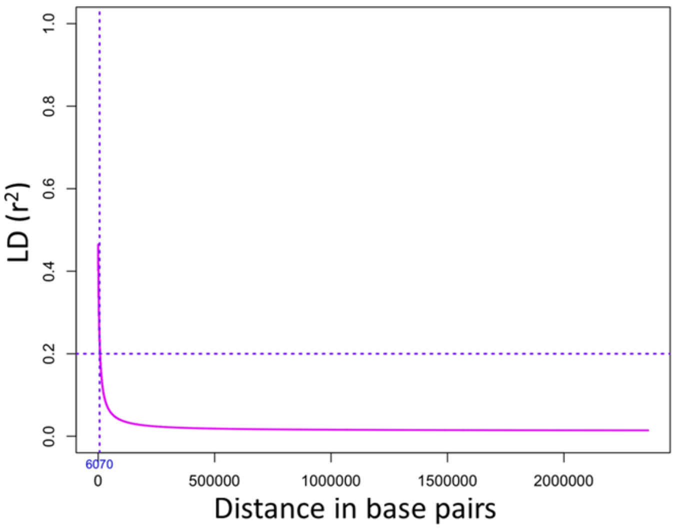


| | LD-decay (in bp) | Total SNPs (per scaffold) |
|---|---|---|
| Scaffold_1 | 4,817 | 236,717 |
| Scaffold_2 | 5,375 | 293,389 |
| Scaffold_3 | 8,283 | 277,886 |
| Scaffold_4 | 4,758 | 298,696 |
| Scaffold_5 | 2,736 | 287,447 |
| Scaffold_6 | 14,760 | 517,295 |
| Scaffold_7 | 4,335 | 301,856 |
| Genome-Wide | 6,070 | 2,213,286 |

**Supplementary Figure 13.** Genome-wide linkage disequilibrium decay plotted against physical distance for MN106-Ref (T_arvense_v2) at an r-squared value of 0.2 and chromosome level LD decay described in the right. Linkage disequilibrium (LD) was calculated using 2,213,286 genome-wide markers with a sliding window of 40 markers.

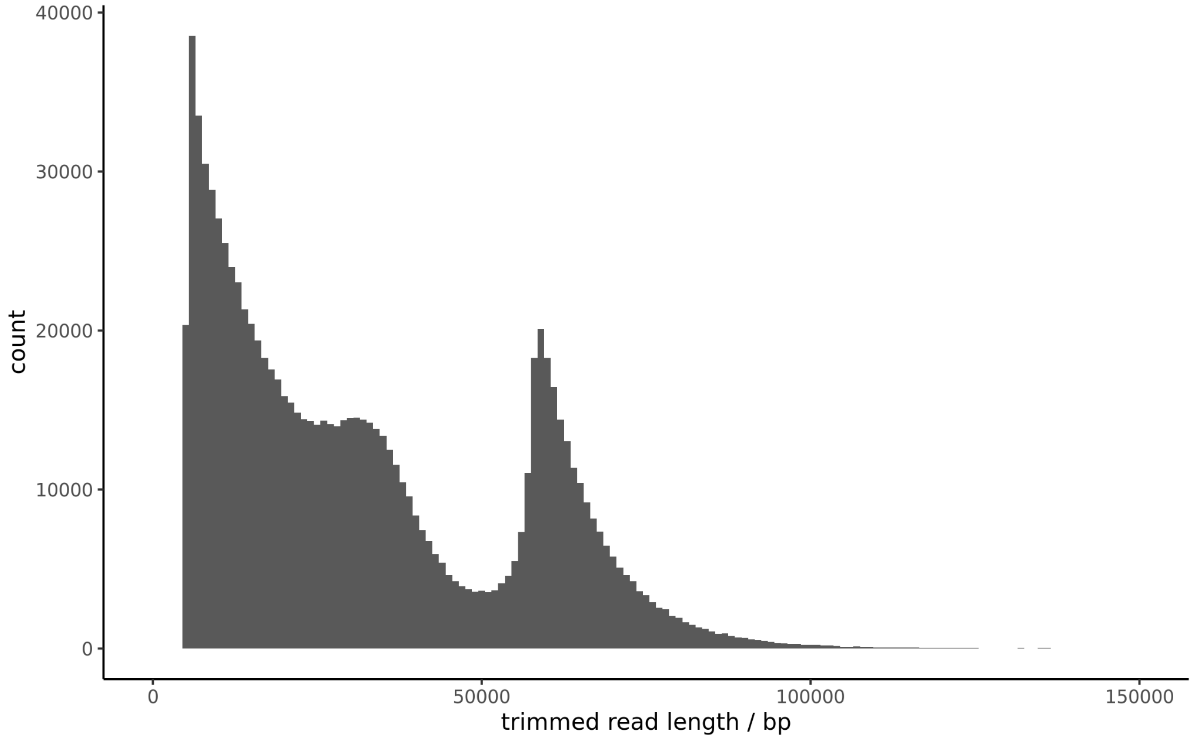


**Supplementary Figure 14.** Read length distribution of trimmed PacBio Sequel II HiFi CLR reads taken forward for assembly with Canu v1.9.

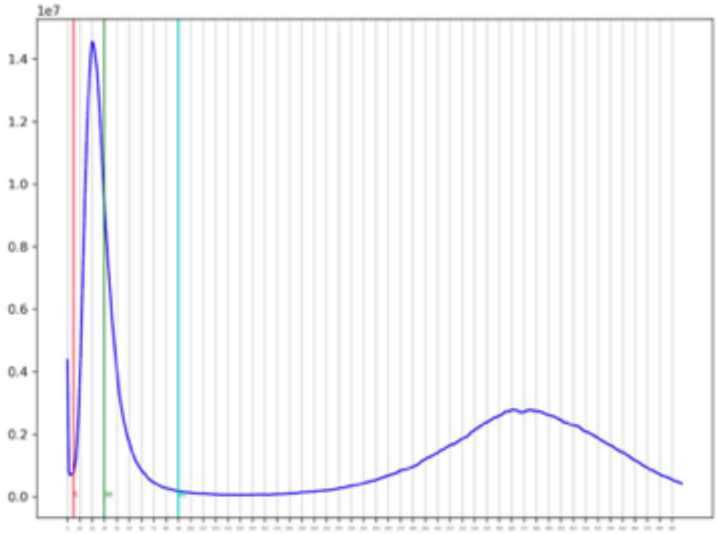


**Supplementary Figure 15.** Distribution of PacBio Sequel II HiFi CLR read mapping depth frequency over assembled contigs, with bimodal peaks due to contig regions with lower depth than the average indicating that they are duplicated.

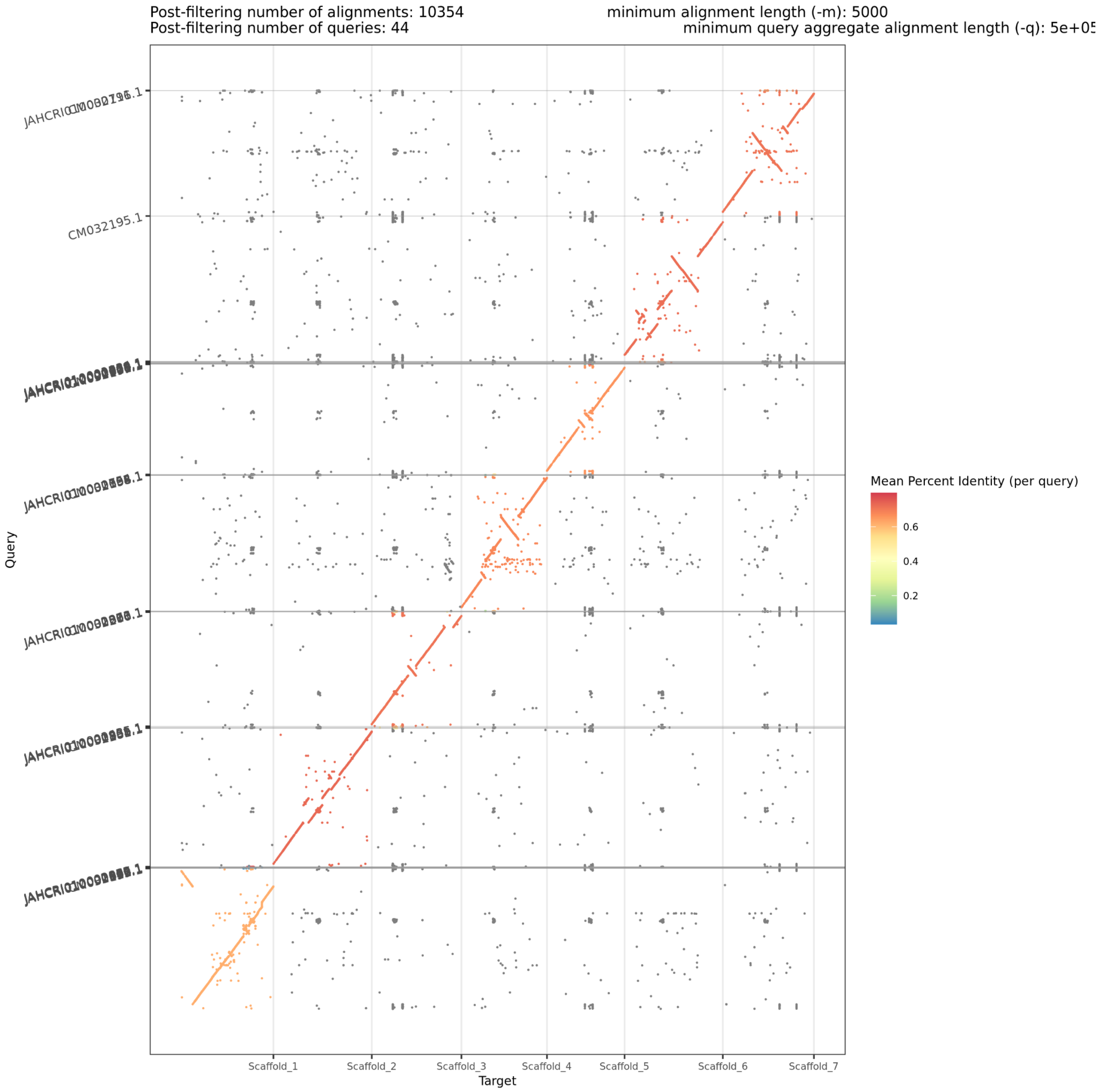


**Supplementary Figure 16.** Synteny between T_arvense_v2 (x-axis) and YUN_Tarv_1.0 (y-axis). Significantly more un-scaffolded contigs from YUN_Tarv_1.0 map to T_arvense_v2 than vice versa, with a total of 44 query sequences from the Chinese accession retained after post-filtering for minimum alignment length 5,000 and aggregate alignment length of 500,000. Most un-scaffolded contigs map to pericentromeric and centromeric regions, which are visible here in addition to the notable mis-scaffolding of centromeric repeats at the telomeric ends of the chromosome-representing scaffolds in YUN_Tarv_1.0.

**Supplementary Table 1.** Estimation of the genome size of *T. arvense* using flow cytometry with *Arabidopsis thaliana*, tomato (*Solanum lycopersicum*), and maize (*Zea mays*) as references.

| **Sample** | **DNA Content (pg)** | **Predicted Genome Size using the reference (Mbp)** |
|---|---|---|
| Field pennycress (MN106-Ref) | 1.09 | NA |
| *Arabidopsis thaliana* | 0.32 | 459 |
| Tomato | 2.05 | 505 |
| Maize | 5.45 | 540 |

**Supplementary Table 2.** Alignment statistics of mRNA-seq reads prior to merging by tissue type.

| sample ID | tissue | replicate | # reads | # alignments | % mapping rate |
|---|---|---|---|---|---|
| 92479 | Cauline leaf | 1 | 26,807,084 | 22,791,298 | 85.02 |
| 92480 | Cauline leaf | 2 | 30,493,022 | 28,502,974 | 93.47 |
| 92468 | Cauline leaf | 3 | 34,703,622 | 32,482,275 | 93.6 |
| 92460 | Green seed | 1 | 34,409,210 | 30,592,247 | 88.91 |
| 92451 | Green seed | 2 | 4,350,126 | 731,192 | 16.81 |
| 92476 | Green seed | 3 | 15,881,224 | 10,421,117 | 65.62 |
| 92454 | Inflorescence | 1 | 37,241,578 | 35,012,072 | 94.01 |
| 92455 | Inflorescence | 2 | 44,709,756 | 42,480,499 | 95.01 |
| 92456 | Inflorescence | 3 | 40,602,958 | 38,486,094 | 94.79 |
| 92471 | Mature seed | 1 | 8,563,360 | 3,050,475 | 35.62 |
| 92459 | Mature seed | 2 | 8,788,058 | 2,824,810 | 32.14 |
| 92475 | Old green silique | 2 | 6,472,836 | 1,831,577 | 28.3 |
| 92467 | Old green silique | 3 | 6,097,296 | 955,425 | 15.67 |
| 92470 | Old green silique | 4 | 8,450,100 | 3,160,559 | 37.4 |
| 92482 | Open flowers | 1 | 27,784,846 | 26,166,091 | 94.17 |
| 92452 | Open flowers | 2 | 48,974,428 | 40,861,167 | 83.43 |
| 92453 | Open flowers | 3 | 33,257,622 | 31,907,812 | 95.94 |
| 92473 | Root 1 week old | 1 | 32,394,826 | 29,934,857 | 92.41 |
| 92474 | Root 1 week old | 2 | 29,742,158 | 27,530,520 | 92.56 |
| 92458 | Root 1 week old | 3 | 33,174,194 | 31,436,791 | 94.76 |
| 92469 | Rosette leaf | 1 | 28,171,012 | 26,329,294 | 93.46 |
| 92478 | Rosette leaf | 2 | 27,902,144 | 26,281,165 | 94.19 |
| 92481 | Rosette leaf | 3 | 31,754,984 | 30,136,304 | 94.9 |
| 92461 | Seed pod | 1 | 39,817,146 | 36,813,142 | 92.46 |
| 92462 | Seed pod | 2 | 37,222,832 | 34,593,888 | 92.94 |
| 92477 | Seed pod | 3 | 27,609,204 | 24,962,517 | 90.41 |
| 92466 | Shoot 1 week old | 1 | 41,735,500 | 38,909,856 | 93.23 |
| 92472 | Shoot 1 week old | 2 | 32,111,696 | 30,143,229 | 93.87 |
| 92457 | Shoot 1 week old | 4 | 35,692,836 | 34,067,138 | 95.45 |
| 92463 | Young green silique | 1 | 37,909,974 | 34,540,606 | 91.11 |
| 92464 | Young green silique | 2 | 36,102,364 | 32,464,393 | 89.92 |
| 92465 | Young green silique | 3 | 35,177,290 | 30,480,559 | 86.65 |

**Supplementary Table 3.** Detailed per-class statistics of the transposable element fraction of the *T. arvense* genome.

| Family | Key Name | Count | bp masked | % masked |
|---|---|---|---|---|
| hAT | DTA | 7,449 | 3,312,483 | 0.63 |
| CACTA | DTC | 12,085 | 6,997,150 | 1.33 |
| Harbinger | DTH | 6,187 | 1,832,186 | 0.35 |
| MuLE | DTM | 18,022 | 8,017,253 | 1.53 |
| Mariner | DTT | 706 | 101,162 | 0.02 |
| Helitron | DHH | 24,151 | 11,129,635 | 2.12 |
| LINE | RIC,RII,RIL,RIX | 26,284 | 11,390,482 | 2.18 |
| Copia | RLC | 37,544 | 31,386,966 | 5.97 |
| Gypsy | RLG | 282,353 | 24,156,3847 | 45.96 |
| LTR | RLA | 9,506 | 5,085,962 | 0.97 |

**Supplementary Table 4**. Description of genes identified in the QTL region (Scaffold_6: 63.85 - 63.95 Mbp) of the BSA analysis of pale seedling phenotype in pennycress.

| Genes within QTL | Gene Description | Mutations (Y/N) |
|---|---|---|
| Tarvense_05935 | Protein of unknown function | N |
| Tarvense_05936 | Similar to DTX25: Protein DETOXIFICATION 25 (*Arabidopsis thaliana* OX=3702) | N |
| Tarvense_05937 | Similar to LHP1: Chromo domain-containing protein LHP1 (*Arabidopsis thaliana* OX=3702) | Y (synonymous) |
| Tarvense_05938 | Protein of unknown function | Y (synonymous) |
| Tarvense_05939 | Similar to At5g24840: tRNA (guanine-N(7)-)-methyltransferase (*Arabidopsis thaliana* OX=3702) | N |
| Tarvense_05940 | Protein of unknown function | Y (Pro59del) |
| Tarvense_05941 | Protein of unknown function | N |
| Tarvense_05942 | Protein of unknown function | N |
| Tarvense_05943 | Similar to AUG7: AUGMIN subunit 7 (*Arabidopsis thaliana* OX=3702) | Y (synonymous) |
| Tarvense_05944 | Protein of unknown function | Y (Asp44Gly) |
| Tarvense_05945 | Similar to At5g17580: BTB/POZ domain-containing protein At5g17580 (*Arabidopsis thaliana* OX=3702) | Y (synonymous) |
| Tarvense_05946 | Similar to BOLA4: Protein BOLA4, chloroplastic/mitochondrial (*Arabidopsis thaliana* OX=3702) | Y (3_prime_UTR) |
| Tarvense_05947 | Similar to PEX19-2: Peroxisome biogenesis protein 19-2 (*Arabidopsis thaliana* OX=3702) | N |
| Tarvense_05948 | Similar to CHAT: (Z)-3-hexen-1-ol acetyltransferase (*Arabidopsis thaliana* OX=3702) | Y (Arg268His) |
| Tarvense_05949 | Protein of unknown function | Y (splice_region_variant) |
| Tarvense_05950 | Similar to MEX1: Maltose excess protein 1, chloroplastic (*Arabidopsis thaliana* OX=3702) | Y (upstream_gene_variant) |
| Tarvense_05951 | Similar to Glycosyl hydrolase 5 family protein (*Chamaecyparis obtusa* OX=13415) | Y (synonymous) |

**Supplementary Table 5.** BUSCO statistics on **a)** initial assembly, immediately after CANU, and **b)** final assembly. Both are derived from orthologs to the *Eudicotyledons odb10* database.

| **a)** C:98.4%[S:74.8%,D:23.6%],F:0.6%,M:1.0%,n2121 | |
|---|---|
| 2086 | Complete BUSCOs (C) |
| 1586 | Complete and single-copy BUSCOs (S) |
| 500 | Complete and duplicated BUSCOs (D) |
| 12 | Fragmented BUSCOs (F) |
| 23 | Missing BUSCOs (M) |
| 2121 | Total BUSCO groups searched |

| **b)** C:98.7%[S:92.1%,D:6.6%],F:0.5%,M:0.8%,n:2121 | |
|---|---|
| 2094 | Complete BUSCOs (C) |
| 1954 | Complete and single-copy BUSCOs (S) |
| 140 | Complete and duplicated BUSCOs (D) |
| 11 | Fragmented BUSCOs (F) |
| 16 | Missing BUSCOs (M) |
| 2121 | Total BUSCO groups searched |

**Supplementary Table 6.** Summary of data provided by each institute and corresponding application.

| Data | Institute | Primary Application |
|---|---|---|
| PacBio Hifi CLR Sequel II | MPI DB | Genome assembly |
| PacBio Hifi CCS Sequel II | University Minnesota | Genome assembly |
| Illumina TruSeq (PCR-free) | GMI | Genome assembly |
| Hi-C sequencing | University Minnesota | Genome assembly |
| Genetic linkage maps | University Minnesota | Genome assembly |
| Bionano maps | University Minnesota | Genome assembly |
| Illumina mRNA-seq (11 tissues) | GMI | Annotation |
| Small RNA libraries (4 tissues) | MPI DB | Annotation |
| Whole genome bisulfite | GMI | DNA Methylation |
| Illumina Novaseq (39 lines) | University Minnesota | Population genomics |
| PacBio Iso-seq | University Minnesota | Transcript variation |
| BSA sequencing | University Minnesota | Population genomics |

**Supplementary Table 7.** Merqury k-mer (k=21) analysis of Illumina HiSeq reads sequenced from the accession in YUN_Tarv_1.0, showing greater QV scores in T_arvense_v2 for the equivalent top 7 scaffolds based on k-mers found uniquely in each assembly and those shared with the read set.

| | **T_arvense_v2** | | | | **YUN_Tarv_1.0** | | | |
|---|---|---|---|---|---|---|---|---|
| | **length** | **uniq. k-mer** | **QV** | **error** | **length** | **uniq. k-mer** | **QV** | **error** |
| Scaffold_1 | 65,519,694 | 1,288,483 | 30.24 | 0.0009 | 73,273,813 | 3,665,265 | 26.13 | 0.0024 |
| Scaffold_2 | 70,024,556 | 1,619,266 | 29.53 | 0.0011 | 72,401,710 | 2,823,932 | 27.23 | 0.0019 |
| Scaffold_3 | 63,812,002 | 969,655 | 31.37 | 0.0007 | 59,618,324 | 2,615,059 | 26.71 | 0.0021 |
| Scaffold_4 | 60,964,055 | 1,833,722 | 28.38 | 0.0015 | 70,763,934 | 2,808,229 | 27.15 | 0.0019 |
| Scaffold_5 | 55,234,666 | 1,400,039 | 29.13 | 0.0012 | 57,454,197 | 3,020,585 | 25.90 | 0.0026 |
| Scaffold_6 | 69,981,056 | 2,370,907 | 27.85 | 0.0016 | 75,772,189 | 4,364,339 | 25.50 | 0.0028 |
| Scaffold_7 | 64,850,309 | 1,826,069 | 28.67 | 0.0014 | 65,366,657 | 2,922,800 | 26.62 | 0.0022 |

**Supplementary Table 8.** Merqury k-mer (k=21) analysis of Illumina HiSeq reads (PCR-free) sequenced from the accession MN106-Ref, showing greater QV scores in T_arvense_v2 for the equivalent top 7 scaffolds based on k-mers found uniquely in each assembly and those shared with the read set.

| | **T_arvense_v2** | | | | **YUN_Tarv_1.0** | | | |
|---|---|---|---|---|---|---|---|---|
| | **length** | **uniq. k-mer** | **QV** | **error** | **length** | **uniq. k-mer** | **QV** | **error** |
| Scaffold_1 | 65,519,694 | 98,643 | 41.44 | 7.2E-05 | 73,273,813 | 4,505,381 | 25.20 | 0.0030 |
| Scaffold_2 | 70,024,556 | 78,174 | 42.74 | 5.3E-05 | 72,401,710 | 4,121,308 | 25.55 | 0.0028 |
| Scaffold_3 | 63,812,002 | 109,566 | 40.87 | 8.2E-05 | 59,618,324 | 3,133,918 | 25.90 | 0.0026 |
| Scaffold_4 | 60,964,055 | 69,169 | 42.67 | 5.4E-05 | 70,763,934 | 4,393,318 | 25.16 | 0.0030 |
| Scaffold_5 | 55,234,666 | 41,707 | 44.44 | 3.6E-05 | 57,454,197 | 4,022,088 | 24.62 | 0.0035 |
| Scaffold_6 | 69,981,056 | 71,047 | 43.15 | 4.8E-05 | 75,772,189 | 5,931,889 | 24.12 | 0.0039 |
| Scaffold_7 | 64,850,309 | 74,777 | 42.60 | 5.5E-05 | 65,366,657 | 4,436,822 | 24.76 | 0.0033 |

**Supplementary Table 9.** Merqury k-mer (k=21) analysis of each total assembly showing relative completeness of k-mers present in each read set from Illumina HiSeq.

| **Assembly** | **Read set** | **k-mers (asm)** | **k-mers (reads)** | **% completeness** |
|---|---|---|---|---|
| YUN_Tarv_1.0 | MN106-Ref | 211,016,166 | 228,654,390 | 92.2861 |
| **T_arvense_v2** | **MN106-Ref** | **227,350,203** | **228,654,390** | **99.4296** |
| both | MN106-Ref | 227,710,775 | 228,654,390 | 99.5873 |
| **YUN_Tarv_1.0** | **SRR14757813** | **223,058,386** | **229,823,560** | **97.0564** |
| T_arvense_v2 | SRR14757813 | 215,415,116 | 229,823,560 | 93.7306 |
| both | SRR14757813 | 227,717,433 | 229,823,560 | 99.0836 |

**Supplementary Information 1**. Manual Curation of predicted miRNAs.

Each miRNA is identified by its Cluster name and genomic location plus mature sequence, below a checkbox of the evaluated features as proposed by Axtell and Meyers (2018) to determine if a putative miRNA can be annotated as such. To evaluate some of the proposed features we used RNA folding form V2.3 (http://www.unafold.org/mfold/applications/rna-folding-form-v2.php) to evaluate the secondary structure of the predicted miRNA precursor loci with default parameters, expect temperature, that we set as 23C, and chose the structure with the lowest ΔG. Homology: predicted mature miRNAs are assigned as homologous from miRbase if it presents a match with two or less nucleotide mismatches. We however made two exceptions based on 100% homology with literature proven mature miRNA: Cluster_8158, which, despite having extra mismatch base in the duplex than allowed, its mature sequence showed a 100% homology with miR408, a well identified miRNA in *Arabidopsis thaliana* (Tatcher *et al.* 2015, Sunkar & Zhu 2004, Abdel-Ghany & Pilon 2008); and Cluster_18198 that also hosts a miRNA with 100% homology with miR396 reported by Soto-Suarez *et al*. 2017. Out of 74 predicted miRNAs, we rejected two loci: Cluster_11184 and Cluster_1133.
